# An electrodiffusive neuron-extracellular-glia model with somatodendritic interactions

**DOI:** 10.1101/2020.07.13.200287

**Authors:** Marte J. Sætra, Gaute T. Einevoll, Geir Halnes

## Abstract

Computational modeling in neuroscience has largely focused on simulating the electrical activity of neurons, while ignoring other components of brain tissue, such as glial cells and the extracellular space. As such, most existing models can not be used to address pathological conditions, such as spreading depression, which involves dramatic changes in ion concentrations, large extracellular potential gradients, and glial buffering processes. We here present the electrodiffusive neuron-extracellular-glia (edNEG) model, which we believe is the first model to combine multicompartmental neuron modeling with an electrodiffusive framework for intra- and extracellular ion concentration dynamics in a local piece of neuro-glial brain tissue. The edNEG model (i) keeps track of all intraneuronal, intraglial, and extracellular ion concentrations and electrical potentials, (ii) accounts for neuronal somatic action potentials, and dendritic calcium spikes, (iii) contains a neuronal and glial homeostatic machinery that gives physiologically realistic ion concentration dynamics, (iv) accounts for electrodiffusive transmembrane, intracellular, and extracellular ionic movements, and (v) accounts for glial and neuronal swelling caused by osmotic transmembrane pressure gradients. We demonstrate that the edNEG model performs realistically as a local and closed system, i.e., that it maintains a steady state for moderate neural activity, but experiences concentration-dependent effects, such as altered firing patterns and homeostatic breakdown, when the activity level becomes too intense. Furthermore, we study the role of glia in making the neuron more tolerable to hyperactive firing and in limiting neuronal swelling. Finally, we discuss how the edNEG model can be integrated with previous spatial continuum models of spreading depression to account for effects of neuronal morphology, action potential generation, and dendritic Ca^2+^ spikes which are currently not included in these models.

**Author summary:** Neurons communicate by electrical signals mediated by the movement of ions across the cell membranes. The ionic flow changes the ion concentrations on both sides of the cell membranes, but most modelers of neurons assume ion concentrations to remain constant. Since the neuronal membrane contains structures called ion pumps and cotransporters that work to maintain close-to baseline ion concentrations, and the brain contains a cell type called astrocytes that contribute in keeping an appropriate ionic environment for neurons, the assumption is justifiable in many scenarios. However, for several pathological conditions, such as epilepsy and spreading depression, the ion concentrations may vary dramatically. To study these scenarios, we need models that account for changes in ion concentrations. In this paper, we present what we call the electrodiffusive neuron-extracellular-glia model (edNEG), which keeps track of all ions in a closed system containing a neuron, the extracellular space surrounding it, and an astrocytic “domain”. The edNEG model ensures a complete and consistent relationship between ion concentrations and charge conservation. We envision that the model can be used to study a range of pathological conditions such as spreading depression and, hence, be of great value for the field of neuroscience.

## Introduction

Computational modeling in neuroscience has largely focused on simulating the electrical activity of neurons and networks of such, while ignoring other components of brain tissue, such as glial cells and the extracellular space. Within that paradigm, biophysically detailed neuron models are typically based on a combination of a Hodgkin-Huxley type formalism for membrane mechanisms (see, e.g., [1, 2]), and cable theory for how signals propagate in dendrites and axons (see, e.g., [3, 4]). Two underlying assumptions in these standard models are that (i) the extracellular space (ECS) is isopotential and grounded, and thus does not affect the neurodynamics, and (ii) that the concentrations of the main charge carriers (Na^+^, K^+^, and Cl^−^) remain constant over the simulated period.

The assumptions (i-ii) are never strictly true. Neuronal activity does give rise to electric fields in the ECS, and in principle, a field will affect the membrane potential dynamics of both the neuron that gave rise to the field and of its neighbors. Such so-called “ephaptic” effects have been the topic of many studies (see, e.g., [5–12]). Furthermore, electrical signals in neurons are generated by transmembrane ion fluxes, which will alter both intra- and extracellular ion concentrations. This may change ionic reversal potentials, and the effect that this may have on neurodynamics has also been the topic of many studies (see, e.g., [13–17]).

As homeostatic mechanisms, such as ion pumps and cotransporters, strive to maintain ion concentrations close to constant baseline levels [18], and as ECS potentials tend to be very small compared to the membrane potentials of neurons [9], the assumptions (i-ii) are still warranted under many conditions. However, there are also many conditions where these assumptions are not justified. For example, in tightly packed bundles of axons, it is likely that the activity in one axon may affect its neighbors both (ephaptically) through the electric field that it evokes [6, 10], and through the ion concentration changes it generates in the narrow ECS separating them [16]. On the much larger spatial scale of brain tissue, spreading depression (SD) and a number of related pathological conditions are associated with dramatic shifts in the K^+^ concentration and giant DC-like voltage gradients in the ECS, which may be as large as several tens of millimolar and millivolts, respectively [19–24]. The pathophysiology of SD is believed to largely depend on the dynamics of extracellular K^+^ [20, 23, 25, 26], which in turn is likely to involve numerous processes such as neuronal re-uptake, electrodiffusion through the ECS, and glial spatial buffering processes [27].

Accurate modeling of conditions that involve notable changes in ion concentrations and ECS potentials requires a unified, electrodiffusive framework that ensures conservation of ions and charge, and a physically consistent relationship between ion concentrations and electrical potentials in both the intra- and extracellular space [17]. Until recently, models that were consistent in this regard (see, e.g., [16, 27–29]), had not accounted for morphological aspects of neurons, such as, e.g., the differential expression of membrane mechanisms in dendrites versus somata. The morphology of a neuron is important not only for its somatodendritic signaling and integration of synaptic inputs but also has implications for the extracellular dynamics of electrical potentials and ion concentrations. For example, the large extracellular shifts seen during SD have been suggested to originate in superficial layers of hippocampus and cortex, and to depend strongly on ion channel openings in the apical dendrites of pyramidal cells [19, 20, 24, 30–32]. Hence, morphological details may have important implications also for understanding dynamical processes at the level of brain tissue.

Recently, we developed the electrodiffusive Pinsky-Rinzel (edPR) model, which we believe is the first model that combines morphologically explicit neuron modeling with biophysically consistent modeling of ion concentrations, electrical charge, and electrical potentials in both the intra- and extracellular space [17]. In that work, we equipped the well established Pinsky-Rinzel model [33] with a homeostatic machinery and equations for ion concentration dynamics in the intra- and extracellular space. The objective was to supply the neuroscience community with a model that can simulate neural dynamics not only under a steady-state scenario (S1), where the homeostatic machinery succeeds in maintaining ion concentrations close to baseline, but also under a scenario (S2) where homeostasis is incomplete, so that ion concentrations change over time.

Two important contributors to ion concentration dynamics were not accounted for in the edPR model, namely the effects of glial cells and cellular swelling or shrinkage. In particular, a type of glial cells called astrocytes is known to be important for regulating the ionic content of the ECS [34], and especially for the uptake of excess K^+^ that may develop during neuronal hyperactivity [35–38]. Furthermore, when ion concentrations change in neurons, astrocytes, and the ECS, it will cause osmotic pressure gradients over the cellular membrane. This can lead to cellular swelling or shrinkage [39–42], which in turn will alter the ionic concentrations in the swollen or shrunken volumes. Cellular swelling and a corresponding shrinkage of the ECS is, for example, and important trademark of pathological conditions such as seizures and SD [18, 23, 43].

In this work, we present an expanded version of the edPR model, which also accounts for effects of glial ion uptake and neuronal and glial swelling due to osmotic pressure gradients. In the expanded version, which we will refer to as the electrodiffusive neuron-extracellular-glia (edNEG) model, the neuron and glial domain interact through a shared ECS. The edNEG model thus includes the main machinery responsible for ion concentration dynamics in a “unit” piece of brain tissue, i.e., corresponding to a single neuron, and the ECS and glial ion uptake that it has to its disposal. The edNEG model has six compartments, two for each of the three domains. It has the functionality that it (1) keeps track of all ion concentrations (Na^+^, K^+^, Ca^2+^, and Cl^−^) in all compartments, (2) keeps track of the electrical potential in all compartments, (3) has different ion channels in neuronal soma and dendrites so that the neuron can fire somatic action potentials (APs) and dendritic calcium spikes, (4) contains the neuronal and glial homeostatic machinery that maintains a realistic dynamics of the membrane potential and ion concentrations, (5) accounts for transmembrane, intracellular and extracellular ionic movements due to both diffusion and electrical migration, and (6) accounts for cellular swelling of neurons and glial cells due to osmotic pressure gradients.

In this first implementation, we study the edNEG model as a closed system, i.e., ions are conserved and confined to stay within the six-compartment system. We focus on illustrating the model tuning required to achieve (1)-(6) and the exploration of its dynamical properties. We show that, like for the edPR model, this closed system (i) has a stable resting state, (ii) maintains steady-state firing for (S1) moderate neural activity, and (iii) experiences homeostatic breakdown (S2), mimicking the onset of SD, once the activity level becomes too high. As the main novelty of the edNEG model, compared to the edPR model, is the glial compartment, we put an emphasis on examining the role of glia in making the neuron more tolerable to hyperactive firing and in limiting neuronal swelling.

## Results

### An electrodiffusive Pinsky-Rinzel model with neuron-glia interactions and cellular swelling

The edNEG model consisted of a neuron, an extracellular (ECS) domain, and a glial domain, all of which had two compartments (Fig 1). For the neuron, the two compartments represent the somatic (bottom) and dendritic (top) parts of its morphology, as in the original Pinsky-Rinzel model [33]. For the ECS, the two compartments represent the average ECS that the single neuron has to its disposal surrounding its somata (bottom) and dendrites (top). Finally, the glial cells most involved in ion homeostasis, the astrocytes, are typically interconnected via gap junctions into a continuous syncytium. The glial compartments could thus be interpreted not as two compartments of a single glial cell, but rather as a representative for the average glial buffering surrounding the neural somata (bottom) and dendrites (top). Like for the ECS, we will hence refer to the glia as a “domain” rather than to a single glial cell.

**Fig 1.**
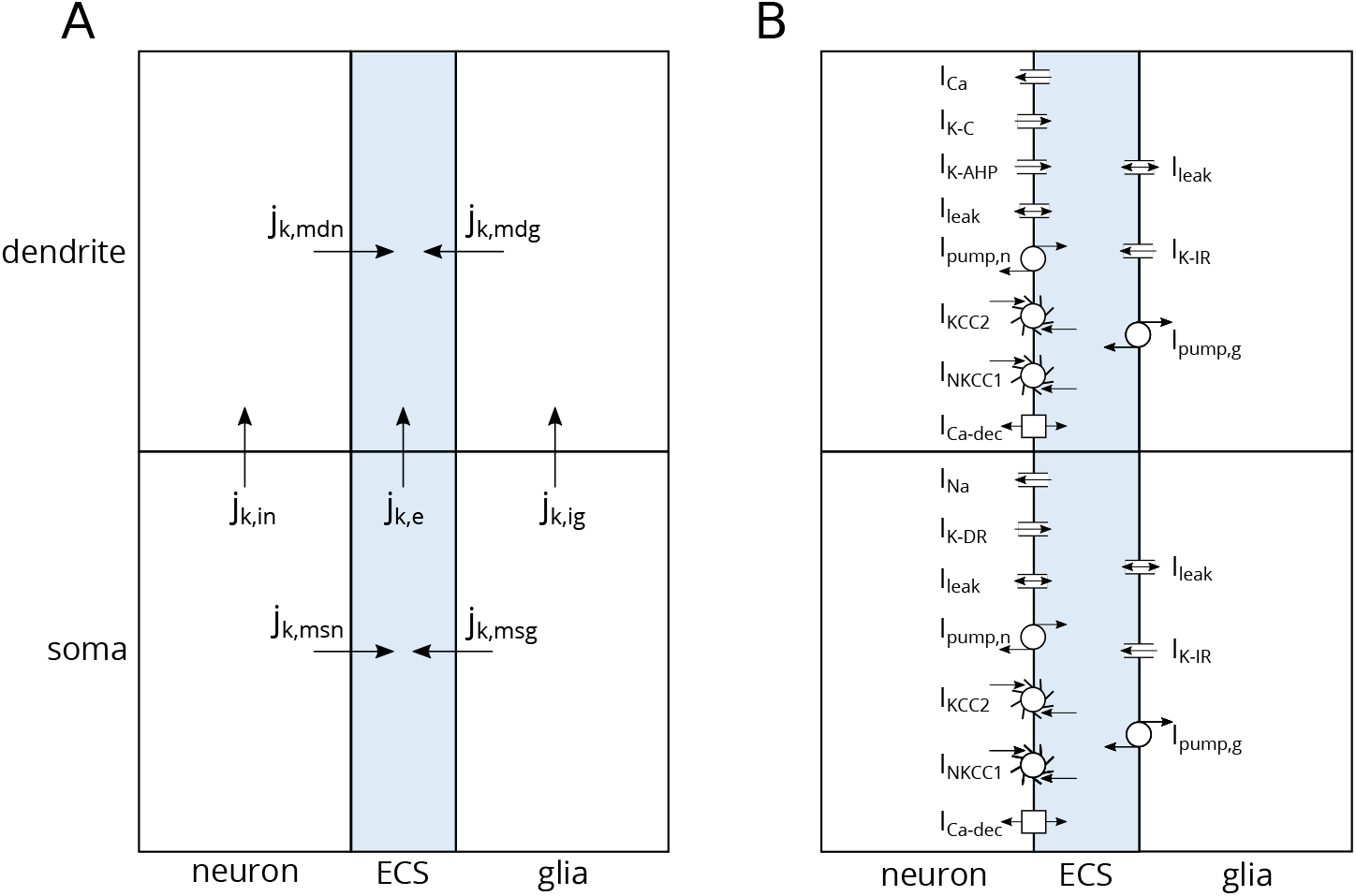
Model architecture. **(A)** The edNEG model contained three domains (neuron, index *n*, ECS, index *e*, and glia, index *g*). Initial neuronal/extracellular/glial volume fractions were 0.4/0.2/0.4. Each domain contained two compartments (soma level, index *s*, and dendrite level, index *d*). Ions of species *k* were carried by two types of fluxes: transmembrane (index *m*) fluxes (*j*_k,msn_, *j*_k,mdn_, *j*_k,msg_, *j*_k,mdg_) and intra-domain fluxes in the neuron (*j*_k,in_), the ECS (*j*_k,e_), and the glial domain (*j*_k,ig_). An electrodiffusive framework was used to calculate ion concentrations and electrical potentials in all compartments. **(B)** The neuronal membrane contained the same mechanisms as in [17]. Active ion channels were taken from [33]. The soma contained Na^+^ and K^+^ delayed rectifier currents (*I*_Na_ and *I*_K−DR_), and the dendrite contained a voltage-dependent Ca^2+^ current (*I*_Ca_), a voltage-dependent K^+^ afterhyperpolarization current (*I*_K−AHP_), and a Ca^2+^-dependent K^+^ current (*I*_K−C_). Both compartments contained Na^+^, K^+^, and Cl^−^ leak currents (*I*_leak_), 3Na^+^/2K^+^ pumps (*I*_pump,n_), K^+^/Cl^−^ cotransporters (*I*_KCC2_), and Na^+^/K^+^/2Cl^−^ cotransporters (*I*_NKCC1_), modeled as in [44]. They also contained Ca^2+^/2Na^+^ exchangers (*I*_Ca−dec_), mimicking the Ca^2+^ decay in [33] and modeled like in [17]. The glial membrane mechanisms were taken from [38], and they were the same in both compartments. They included Na^+^ and Cl^−^ leak currents (*I*_leak_), inward rectifying K^+^ currents (*I*_K−IR_), and 3Na^+^/2K^+^ pumps (*I*_pump,g_).

The neuron and the ECS domain were adopted from the previously published electrodiffusive Pinsky-Rinzel (edPR) model [17] and modified slightly (see Methods). The neuron was based on the Pinsky-Rinzel model, which, despite having only two compartments, can reproduce a variety of realistic firing patterns when responding to somatic or dendritic stimuli, including somatic APs and dendritic calcium spikes [33]. The model for the glial domain was taken from a previous model for astrocytic spatial buffering [38] and added to the edPR model so that both the neuron and glial domain interacted with the ECS. Unlike the previous neuron [17] and glial [38] models that it was based upon, the edNEG model was constructed so that it also accounted for cellular swelling due to osmotic pressure gradients. We implemented the edNEG model using the electrodiffusive KNP framework [17, 38], which consistently outputs the voltage- and ion concentration dynamics in all compartments.

The edNEG model is depicted in Fig 1. Both the neuron and glial domain contained cell-specific and ion-specific passive leakage channels, cotransporters, and ion pumps that ensured a homeostatic ion balance in the system. The neuron contained additional active ion channels that were different in the somatic versus dendritic compartment, making it susceptible to fire somatic action potentials and dendritic Ca^2+^ spikes. Both glial compartments contained inward rectifying K^+^ channels. All included membrane mechanisms are summarized in Fig 1 and described in further detail in the Methods section.

### Action potential firing and resting state in the edNEG model

The neuron in the edNEG model was based on the previous two-compartment Pinsky-Rinzel model for a hippocampal pyramidal neuron in CA3. A feature of the original Pinsky-Rinzel model was that it produced somatic action potentials and dendritic Ca^2+^ spikes. Also, for weak coupling (high intracellular resistance) between the soma and dendrite, the interplay between somatic action potentials and dendritic Ca^2+^ spikes could give rise to a wobbly spike shape (Fig 2A), while for a stronger coupling (low intracellular resistance), the interplay rather lead to a broadening of the AP shape (Fig 2B). These features of the original Pinsky-Rinzel model has been analyzed thoroughly in previous studies [17, 33, 45].

**Fig 2.**
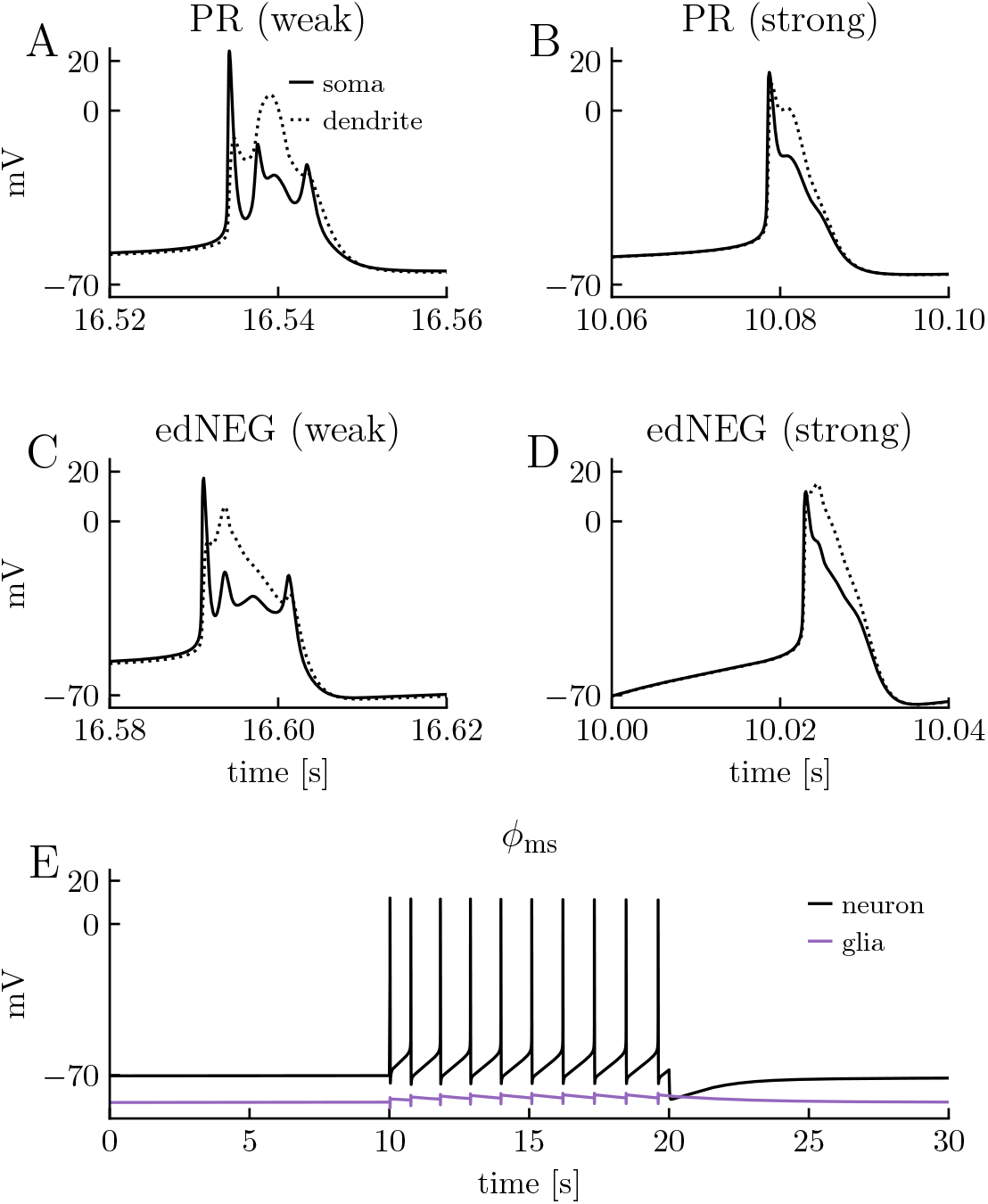
Membrane potential dynamics and resting state in the edNEG model. The neuron in the edNEG model **(C,D)** and the original Pinsky-Rinzel (PR) model **(A,B)** exhibited the same spike shape characteristics for weak coupling **(A,C)** and strong coupling **(C,D)** between the soma and dendrite layers. **(E)** The somatic membrane potential *ϕ*_ms_ of the neuron (black line) and the glial domain (purple line). The neuron received a step current injection to the somatic compartment between *t* = 10 s and *t* = 20 s tuned to give it a firing frequency of 1 Hz. The neuron and the glial domain rested at approximately −70 mV and −83 mV, respectively, when the neuron was not stimulated, and returned to these values after stimulus offset. The stimulus current was 1.35 *μ*A/cm^2^ in **(A)**, 0.78 *μ*A/cm^2^ in **(B)**, 44 pA in **(C)**, and 36 pA in **(D,E)**. **(A,B)** The original PR model was simulated with the code provided in [17]. The coupling conductance of the PR model was 2.26 mS/cm^2^ in **(A)**, and 8.86 mS/cm^2^ in **(B) (C-E)** See methods subsection titled Model tuning for definition of weak and strong coupling in the edNEG model. The strong coupling was used in **(E)**, and as default in all simulations in the reminder of this paper.

To verify that we preserved the characteristic firing properties of the Pinsky-Rinzel model when we made it ion conserving and embedded it within the edNEG model, we implemented two versions of the edNEG model, one with a strong coupling between the soma and dendrite layers, and one with a weak coupling (see Methods for definition of weak and strong coupling). When we stimulated the two versions with constant current injections to the neuronal soma, they elicited spikes that were similar to that of the original Pinsky-Rinzel model: Compare Fig 2A and Fig 2C for weak coupling, and Fig 2B and Fig 2D for strong coupling. Hence, the neuron in the edNEG model preserved the key dynamical properties of the previously developed CA3 hippocampal cell model [33]. The model version with the strong coupling between layers was used as default in all other simulations in this paper.

When we in the edNEG model combined a previous neuron model [17] and a previous glial model [38], and made them share ECS, the original resting state of both the previous models were disturbed. To obtain realistic resting potentials in the new system, we had to re-tune selected parameters (see section titled Model tuning for details). The existence of a realistic resting state for the tuned edNEG model is verified in Fig 2E. It shows a simulation where the neuron was stimulated between *t* = 10 s and *t* = 20 s with a constant current injection that made it fire at 1 Hz. Both the neuron and glial domain stayed at their resting potentials of approximately −70 mV and −83 mV, respectively, when unstimulated (*t* < 10 s), and returned to this resting state after the stimulus had been turned off (*t* > 20 s). The dynamics of the membrane potentials and ion concentrations during on-going activity is analyzed in further detail in the next section.

### Steady-state firing in the edNEG model

In standard (Hodgkin-Huxley type) neuron models, which the original Pinsky-Rinzel model [33] is an example of, the key dynamical variable is the membrane potential. In addition to modeling the membrane potential, the edNEG model presented here keeps track of all neuronal, glial, and extracellular ion concentrations, and accounts for changes in cellular and extracellular volume fractions due to osmotic gradients. It also accounts for the effect that changes in these variables may have on neuronal firing properties.

When the neuron is active, the exchange of ions due to AP firing will be counteracted by the homeostatic mechanisms striving to restore baseline concentration gradients. Hence, we expect that for moderately low neuronal firing, the edNEG model will enter a dynamic steady-state scenario (S1) where homeostasis is successful, and firing can prevail for an arbitrarily long period of time without ion concentrations diverging far off from baseline. We also expect that for a too-high neuronal activity level, the edNEG model will enter a scenario (S2) where the homeostatic mechanisms fail to keep up, and where gradual changes in ion concentrations will lead to gradual changes in neuronal firing properties, and eventually to ceased AP firing.

The existence of a dynamic steady-state scenario (S1) is illustrated in Fig 3, which shows how selected variables vary in the edNEG model during a 1400 s simulation. The neuron received a stimulus from *t* = 1 s to *t* = 600 s that made it fire at 1 Hz (Fig 3A). To examine the steady-state scenario (S1), we have divided the simulation into four phases: an *initial phase* (the first column in Fig 3B-D), covering transient dynamics immediately after stimulus onset), a *steady-state phase* (the second column in Fig 3B-D), covering the last ten seconds of firing, a *recovery phase* (the third column in Fig 3B-D), covering the transient dynamics immediately after the stimulus offset, and a *recovered phase* (the fourth column in Fig 3B-D), covering the last 10 s of the simulation, when the system had returned to the original resting state. In all these phases, we examined the temporal development of the neuronal (Fig 3B) and glial (Fig 3C) reversal potentials, and the neuronal and glial swelling (Fig 3D).

**Fig 3.**
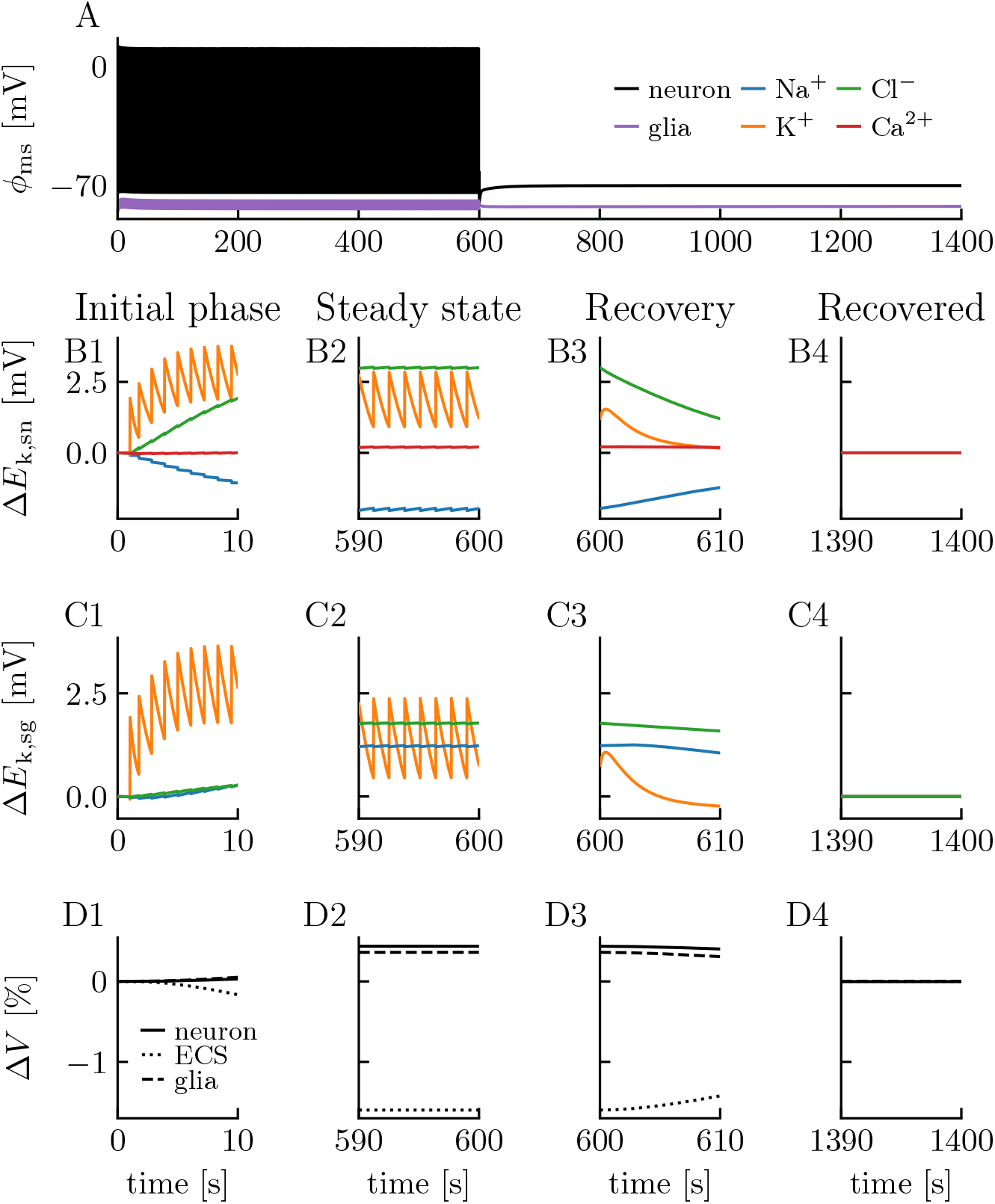
Steady-state firing in the edNEG model. Model response to a 36 pA step-current injection to the somatic compartment of the neuron between *t* = 1 s and *t* = 600 s. The neuron responded with a firing rate of 1 Hz. The simulation covered 1400 s of biological time, and the last 800 s shows recovery to baseline. **(A)** The somatic membrane potential *ϕ*_ms_ of the neuron (black line) and the glial domain (purple line). **(B)** Reversal potential dynamics of the neuronal soma for all ion species (Na^+^, K^+^, Cl^−^, Ca^2+^) shown in terms of their deviance from baseline values. **(C)** Reversal potential dynamics of the glial “soma” for ion species *k* (Na^+^, K^+^, Cl^−^). The glial domain did not contain any Ca^2+^ channels. **(D)** Volume dynamics of the three domains shown in terms of relative changes. Volume changes were computed for the whole domain (soma layer + dendrite layer). Initial neuronal/extracellular/glial volume fractions were 0.4/0.2/0.4. **(B-D)** Rows 1-4 show four selected time intervals, (1) initially after stimulus onset, (2) when the system had reached dynamic steady state, (3) initially after stimulus offset, and (4) when the system had restored baseline.

In the initial phase, the concentrations of all ion species varied with time due to the influxes and effluxes associated with cellular activity. In Fig 3B-C, the concentration variations in the soma layer are reflected in the ionic reversal potentials (*E*_k_), which are proportional to the logarithm of the ratio between the extra- and intracellular ion concentration of a given species *k* (cf. Eq 32). As the soma contained no Ca^2+^ channels, variations in *E*_Ca_ were very small, although not strictly zero, since minor concentration shifts could occur due to electrodiffusion of Ca^2+^ between the soma and dendrite layer. The glial domain did not contain any Ca^2+^ conducting channels. For the other ion species, *E*_k_ had a zig-zagging shape, most pronounced for *E*_K_, where the upstroke reflects the efflux of K^+^ during the repolarization phase of an action potential (AP), while the downstroke reflects the homeostatic mechanisms that were active between APs, working to restore the baseline concentrations. The fact that *E*_K_ showed the largest variations was as expected, as the extracellular K^+^ had the lowest baseline value of all ion species, and therefore experienced the largest relative changes during AP firing.

The homeostatic recovery between APs was incomplete during the initial phase, and the reversal potentials zig-zagged away from baseline for each consecutive AP (Fig 3B1,C1). However, the gradual divergence from baseline increased the homeostatic activity, so that after a period of regular firing, the system entered a dynamic steady-state phase where the zigs and the zags became equal in magnitude, and the reversal potential did not deviate further from baseline (Fig 3B2,C2).

In this simulation, *E*_K_ deviated by maximally ~3 mV from the baseline reversal potential, which was not enough to have a visible impact on the regular firing of the neuron. Hence, the edNEG model supported a steady-state scenario (S1), where the neuron could fire regularly and continuously without dissipating its concentration gradients. For firing in scenario S1, the neuron in the edNEG model performs similarly to the original Pinsky-Rinzel model [33], which does not model ion concentrations, but assumes that they remain constant.

When the stimulus was turned off, the recovery phase started. The membrane potentials returned rapidly to values very close to the resting potential (Fig 3A), while the ion concentrations (and thus the reversal potentials) returned more slowly towards baseline (Figs 3B3,C3). At the end of the simulation, ion concentrations had recovered the baseline values (Figs 3B4,C4). If we define recovery (rather arbitrary) as the time it took for all reversal potentials to return to values less than 0.1 mV away from their resting baseline values, recovery took about 300 s, i.e., it occurred at about *t* = 900 s. The fact that the membrane potentials were almost constant during the recovery of the reversal potentials, indicates that the ion concentration recovery was due to a close-to electroneutral exchange of ions over the neuronal and glial membranes. Hence, the edNEG model predicts that “memories” of previous spiking history may linger in a neuron for several minutes, in the form of altered concentrations, even if it appears to have returned to baseline by judging from its membrane potential.

When the ion concentrations changed, so did the osmotic pressure gradients. This caused the neuron and glial domain to swell over the simulated time course (Fig 3D). Given the rather modest concentration changes observed during 1 Hz firing, the cellular swelling was not dramatic. The neuron swoll maximally by 0.44 %, the glial domain by maximally 0.36 %, and the extracellular space shrunk correspondingly by 1.60 % (Fig 3D2). As initial neuronal/extracellular/glial volume fractions were 0.4/0.2/0.4, this preserved the total volume. After the stimulus was turned off, the three domains recovered their original volume fractions (Fig 3D3-D4), and at about *t* = 1100 s, all volumes were less than 0.01 % away from baseline.

### Homeostatic breakdown in the edNEG model

The existence of a scenario (S2) where the homeostatic mechanisms fail to keep up with the neuronal exchange is illustrated in Fig 4. There, the neuron received a strong input current (150 pA) for three seconds, which gave it a high firing rate (Fig 4A1). While the neuron fired, ion concentrations gradually changed, leading to changes in ionic reversal potentials (Fig 4B1,C1), which in turn caused a gradual depolarization of the neuron and made it fire even faster. The neuron could tolerate this strong input for only a little more than 2 s before it became unable to re-polarize to levels below the AP firing threshold, and the firing ceased due to a permanent inactivation of the AP generating Na^+^ channels. This condition, when a neuron is depolarized to voltage levels making it incapable of eliciting further APs, is known as *depolarization block*. It is a well-studied phenomenon, often caused by high extracellular K^+^ concentrations [46]. This kind of dynamics can not be captured with standard neuron models constructed under the assumption that ion concentrations remain constant under the simulated period. The dynamical characteristics of the neuron in the S2 scenario resembled what we saw in a previous study, including only the neuron and the ECS [17], and is not further analyzed here.

**Fig 4.**
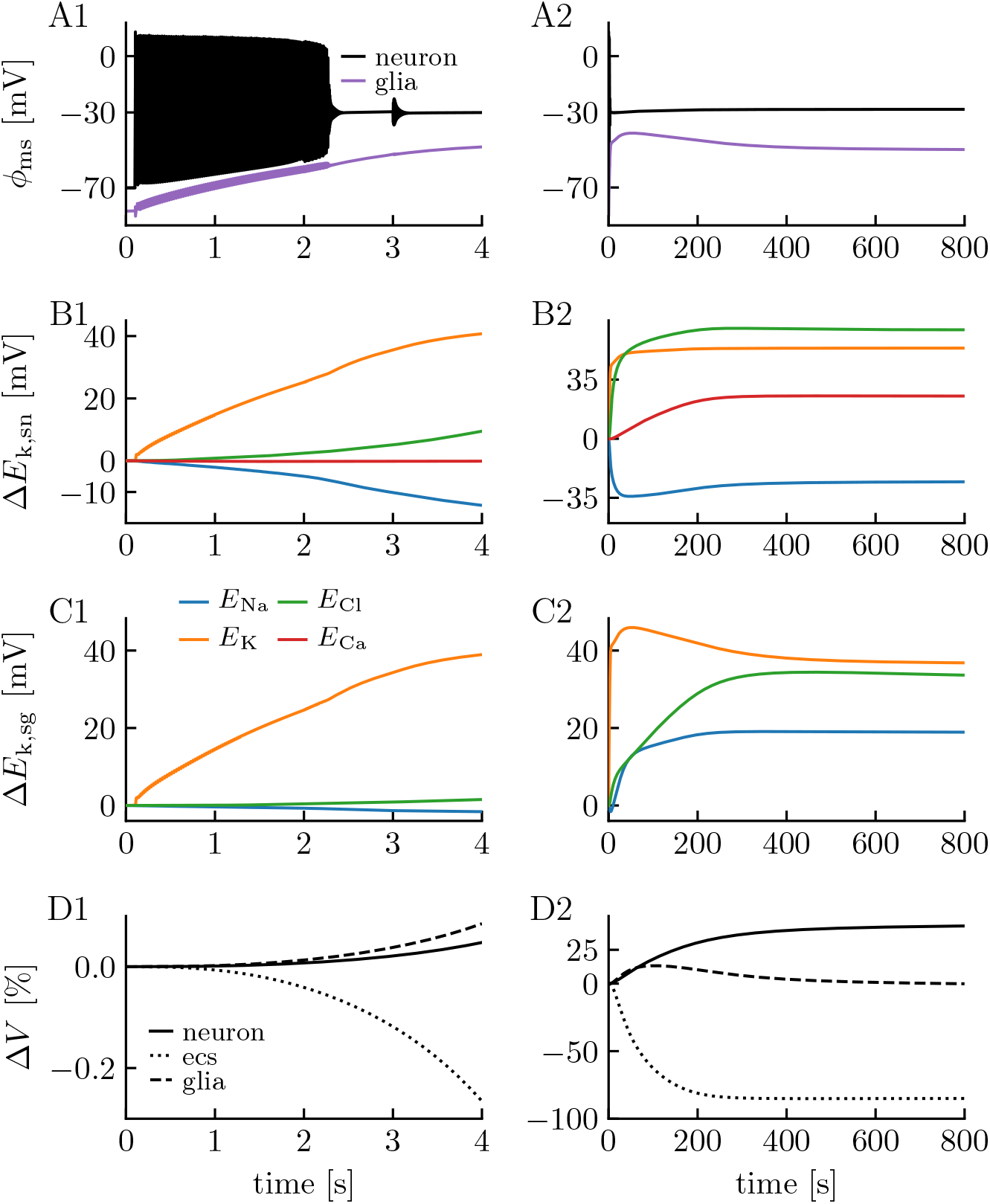
Homeostatic breakdown in the edNEG model. Model response to a 150 pA step-current injection to the somatic compartment of the neuron between *t* = 0.1 s and *t* = 3 s. **(A1)** The neuron responded with an initial firing rate of 50 Hz, but both the firing rate and spike shapes varied throughout the simulation due to variations in the ion concentrations. Both the neuron and glial domain experienced a gradual depolarization throughout the simulation, and the neuron eventually went into depolarization block. The gradually changing dynamics patterns were due to **(B-C)**. **(D)** The system experienced massive neuronal and glial swelling. The first row **(A1-D1)** shows dynamics on a short time scale, and the second row **(A2-D2)** shows the same simulation on a longer time scale.

We note that although the input was turned off after 3 s, the neuron lingered in depolarization block, and continued to dissipate its concentration gradients so that changes in ionic reversal potentials and cellular swelling went on for a long time (Fig 4B2,C2,D2). Interestingly, the glial swelling was transient. After having swollen by 13.2 % during the first 100 s of the simulation, it began to shrink, while the neuron swoll monotonously during the entire simulation. At the end of the simulation, ionic reversal potentials were several tens of millivolts away from their baseline values, the neuron had swollen by 42.6 %, and the glial and ECS domains had shrunken by 0.13 % and 85.0 %, respectively.

In this simulation, the system never returned to its baseline resting state, and the neuron never regained its ability to elicit APs. This has previously been referred to as a *wave-of-death-like* dynamics [44, 47]. It also resembles the neural dynamics seen under the onset of SD [44], but during SD, neurons tend to recover baseline activity after about one minute as the SD wave passes [23]. Putatively, this recovery depends on K^+^ being transported away from the local region by ECS electrodiffusion and spatial buffering through the astrocytic network, and quite likely also vascular clearance. As the edNEG model studied here represented a local and closed system, such spatial riddance of K^+^ did not occur, but we anticipate that recovery might be observed if the edNEG model were expanded to a spatially continuous model (see Discussion for more on this).

### Soma versus dendrite

It is known that the leading edge of the SD wave tends to occur in the layers containing the apical dendrites [20, 32]. Inspired from this, we wanted to explore if the edNEG model expressed such layer-specific differences. Assuming that the SD wavefront coincides with neurons going into depolarization block, we used the simulation from Fig 4 for this comparison. We studied the ECS K^+^ concentration and neuronal swelling in the soma and dendrite layer at a short time scale, as the neuron approached depolarization block (Fig 5A1), and at a longer time scale, when the neuron lingered in depolarization block (Fig 5A2).

**Fig 5.**
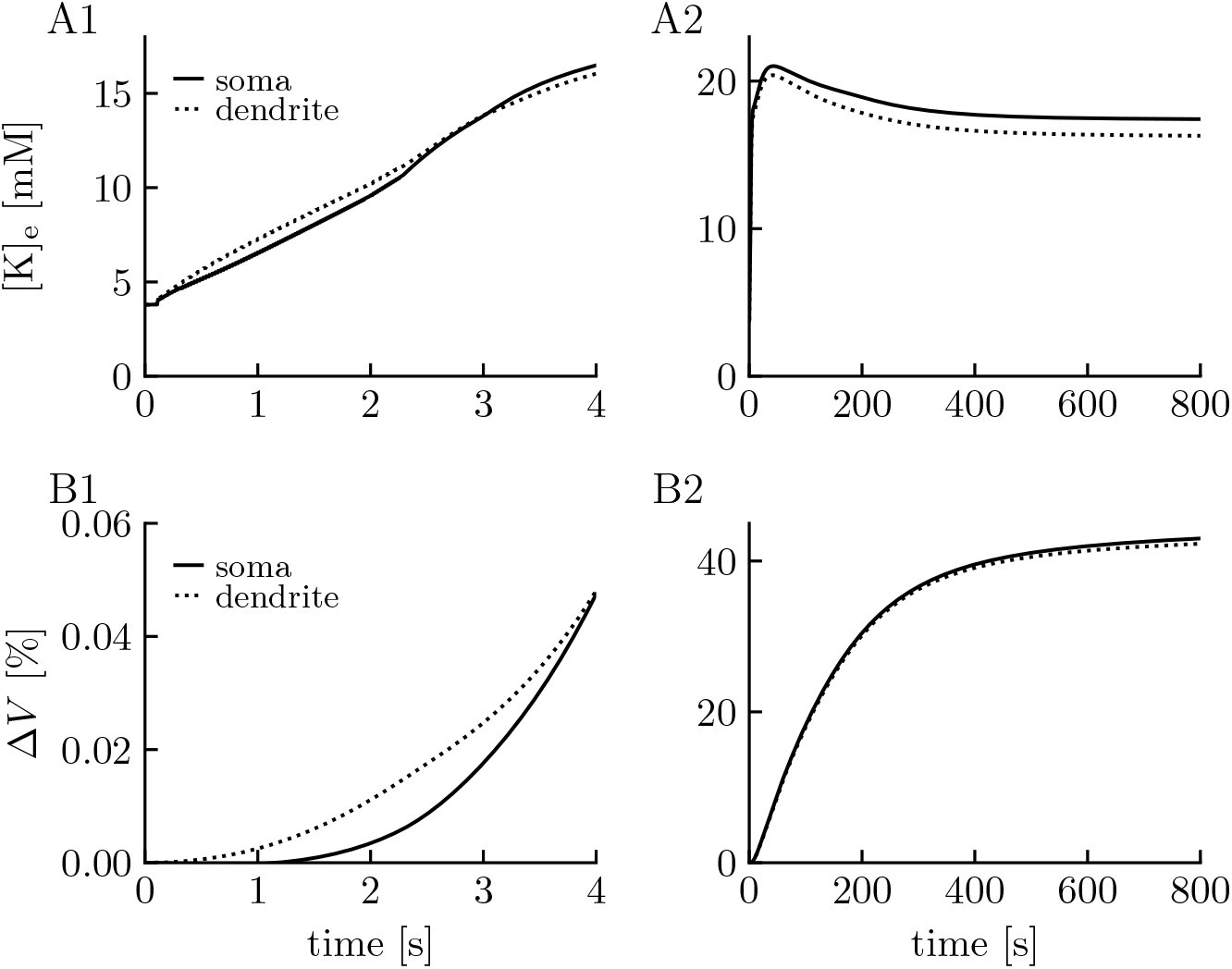
Extracellular potassium and neuronal swelling in the soma versus dendrite layer. Comparison of **(A)** extracellular K^+^ dynamics, and **(B)** neuronal swelling in the soma versus dendrite layer for the simulation in Fig 4. Panels **(A1,B1)** show the first 4 s of the simulation, where the neuron went into depolarization block around *t* = 2.3 s, while **(A2,B2)** show the long term effect of the depolarization block. In the results shown, we stimulated the soma with an inward K^+^ current, but the results were very similar when we stimulated with an inward Na^+^ current, or when the stimulus was applied to the dendritic compartment instead of the soma.

In line with the notion that the dendritic layer is the leading edge of the SD wavefront, Fig 5A1 shows that the ECS K^+^ concentration during neuronal firing (*t* < 2.3 s) was highest in the dendrite layer, but was bypassed by the ECS K^+^ concentration in the soma layer shortly after the neuron had entered depolarization block, after which it stayed highest in the soma layer (Fig 5A2). Similarly, the neuronal dendrite also swoll more than the soma during neuronal firing (Fig 5B1), whereas the somatic swelling caught up after the neuron had entered depolarization block (Fig 5B2). To some degree, these observations are in agreement with the notion that SD initiates in dendritic layers, although the differences between the layers were admittedly rather small in the edNEG model (see Discussion for further comments).

### The effect of the glial domain on neuronal tolerance levels

As the subsystem containing only the neuron and ECS was studied thoroughly in [17], we here put an emphasis on exploring what difference the new glial domain made for the system, especially in terms of how it (i) affected the neuronal tolerance level for AP firing, and (ii) how it affected the dynamics of ECS K^+^ concentrations and cellular swelling. To do this, we compared two versions of the model, one being the full edNEG model *with* glia included, and one *without* glia included. In the latter case, we removed the glial influence by setting all glial membrane conductances and water permeabilities to zero, i.e., we sealed the glial membrane.

When comparing, we wanted to make sure that the neuron fired with the same rate in both versions, something that we could not control using a continuous step current injection, partly because the two model versions had a different response to the same stimulus, and partly because the firing rate could vary over time as the ion activity-induced changes in ionic reversal potentials concentrations changed. To control the firing rate, we, therefore, used a stimulus protocol where we stimulated the neuronal soma periodically with a train of brief and strong 10 ms pulses, each evoking a single AP.

Both versions of the model could maintain sustained regular firing (cf. scenario S1 in Fig 3), provided that the stimulus frequency was low enough, as in the example in Fig 6A,C where the input pulse frequency (and resulting firing rate) was 4 Hz. For the 4 Hz simulations, we verified that both model versions could sustain regular firing for at least 1000 s. However, all other simulations considered in Fig 6 were run for only 90 s, so that “sustained” in this context will mean “sustained for at least 90 s”. We chose to stop the simulations at *t* = 90 s, partly to reduce computation time, and partly because this is a typical time window within which an SD wave passes [23].

**Fig 6.**
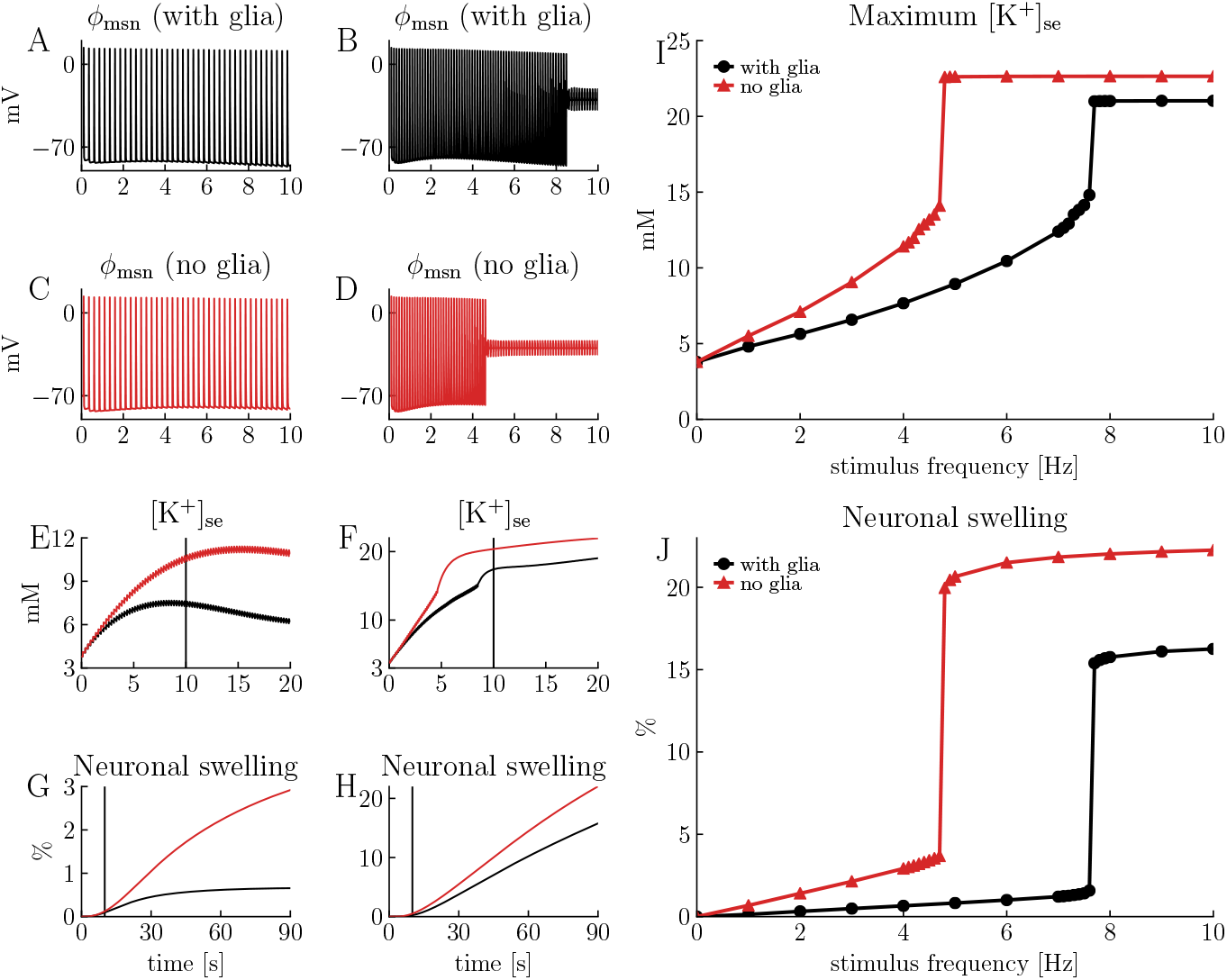
Neuronal firing with and without the presence of glia. The edNEG model tolerated higher neuronal firing frequencies when glia was present. Simulations show responses to trains of 10 ms step-pulses of 320 pA injected to the neuronal soma, each inducing exactly one action potential. The first pulse was applied at *t* = 0.1 s and all simulations covered 90 s of biological time. **(A,C)** Somatic membrane potential of the neuron responding to a pulse train with frequency of 4 Hz, in the case when glia was present **(A)** and not present **(C)**. **(B,D)** Somatic membrane potential of the neuron responding to a pulse train with frequency of 8 Hz, in the case when glia was present **(B)** and not present **(D)**. **(E)** ECS K^+^ concentrations (in soma layer) during the simulations in **(A)** (black line) and **C** (red line). **(F)** ECS K^+^ concentrations (in soma layer) during the simulations in **(B)** (black line) and **(D)**(red line). **(I-J)** Summary of 20 simulations (as those in **(A-D)**) with varying stimulus frequency, showing **(I)** maximum extracellular (soma) K^+^ concentration as a function of stimulus frequency, and **(I)** relative neuronal volume change at the end of the simulations (*t* = 90 s) as a function of stimulus frequency. **(A-D)** show the 10 first seconds of the simulations, and **(E-F)** show only the 20 first seconds of the simulations.

Whereas the membrane potential dynamics at 4 Hz firing were very similar in the versions with and without glia, the ion concentration dynamics were not (Fig 6E). In the case without glia, the ECS K^+^ concentration peaked at 11.4 mM, while it stayed below 7.7 mM when glia was included. In both versions, however, the K^+^ concentration reached a ceiling level and eventually stabilized at a constant value, so that the system entered a dynamic steady state (cf. scenario S1 in Fig 3).

To study homeostatic breakdown (cf. scenario S2 in Fig 4), we increased the stimulus frequency until neither the version with, nor the version without, glia could maintain sustained regular firing for 90 s, as in the example in Fig 6B,D where the input pulse frequency (and resulting initial firing rate) was 8 Hz (Fig 6A,C). Both versions of the model then sustained 8 Hz firing for only a limited period. In both versions, the 8 Hz firing caused the ECS K^+^ concentration to increase (Fig 6F) until it exceeded a ceiling level where the neuron entered depolarization block and AP firing ceased.

As the increase in the ECS K^+^ concentration was much faster when glia was not present, the version without glia entered depolarization block after less than 5 s of activity (Fig 6D), while the version including glia maintained the 8 Hz AP firing for almost 9 s (Fig 6B). The time points where depolarization block was reached can be seen as a dent in the ECS K^+^ concentration curves (Fig 6F), which occurred at a concentration of about 14 mM in the version without glia and at about 15 mM in the version with glia included.

We were surprised to observe that the afterhyperpolarization following APs increased during the journey towards depolarization block in Fig 6B, despite the K^+^ reversal potential becoming more depolarized during the simulation. When exploring this phenomenon, we found that the increased afterhyperpolarization was caused by the electrogenic 3Na^+^/2K^+^ pump, which increased its activity level as the intracellular Na^+^ concentration and ECS K^+^ concentration increased during the simulation (cf. Eq 62), leading to an increased outward current. Such hyperpolarization by the ATP-ase driven 3Na^+^/2K^+^ pump has also been reported in other studies [48, 49].

Fig 6I summarizes a number of simulations of the model versions with and without glia, and shows the peak ECS K^+^ concentration (maximum reached during simulation) as a function of stimulus frequency. The highest tolerated frequency, i.e., the maximum stimulus frequency for which the neuron could sustain AP firing throughout the simulated 90 s, is easily identifiable as the point where the curves make a sharp dent. With glia present, the edNEG model could sustain firing up to 7.6 Hz (black curve), while without glia, it could only maintain regular firing for frequencies up to 4.7 Hz (red curve). Also, the presence of glia reduced the peak ECS K^+^ concentration occurring after homeostatic breakdown from about 23 mM (black curve) to about 21 mM (red curve). The differences were due to the glial support in clearing the ECS from excess K^+^.

As the ion concentrations changed during the simulations, so did the osmotic pressure gradients over the membrane, and this caused cellular swelling and ECS shrinkage. The swelling depended not only on K^+^, but on all ion concentrations changing during the simulation. However, the pattern of how the presence of glia affected the swelling of the neuron was similar to how it affected the ECS K^+^ concentration (Fig 6G,H,J). During steady-state firing, the neuron swoll by up to only 1.6 % when glia was present, and by 3.7 % when glia was not present (Fig 6J for frequencies below the “dent”-frequency).

The swelling was much more dramatic in the simulations where the neuron entered depolarization block. For the maximal stimulus frequency (*f* = 10 Hz), the neuron had swollen by 16.3 % when glia was present, the glial domain had swollen by 13.1 %, and the ECS had shrunken by 58.8 % at the end of the simulation (glial and ECS volume fractions were not included in the plot, but were computed in the same simulation). We note again that the simulation ended at *t* = 90 s, and the glial swelling of 13.1 % was close to the peak glial swelling (13.2 %) seen at *t* = 100 s in Fig 4. When glia was not present, neural swelling was more dramatic, and the neuron had then swollen by 22.3 % at the end of the simulation, with a corresponding shrinkage of the ECS by 44.5 %.

## Discussion

We presented the edNEG model for local ion concentration dynamics in brain tissue containing a neuronal, extracellular, and glial domain (Fig 1). The model contained essential ion channels and homeostatic mechanisms, and accounted for somatodendritic signaling by neurons, for electrodiffusive ion concentration dynamics within all domains, as well as for neuronal and glial swelling due to concentration-dependent osmotic pressure gradients.

We demonstrated that the edNEG model had realistic dynamical properties in the sense that it supported a scenario (S1) when the homeostatic mechanisms could maintain constant ion concentrations so that the neuron could maintain low firing frequencies for an arbitrary long time (Fig 3), and a scenario (S2) when the neuron fired too fast for the homeostatic mechanisms to keep up, so that ionic concentrations gradually changed, leading eventually to the neuron entering depolarization block and losing its ability to generate further action potentials (Fig 4). The first scenario (S1) represents normal physiological conditions and could be modeled fairly well with simpler and more conventional neuronal models assuming constant ion concentrations and reversal potentials. The second scenario (S2) resembles the onset of pathological conditions such as spreading depression (SD) and the wave of death [44] and requires models that explicitly account for variations in ion concentrations. Of course, concentration effects are not only relevant during homeostatic breakdown. As Fig 3 showed, concentrations vary at a much slower time course than the membrane potential, and may give neurons “memories” of previous spiking history that may last for several minutes. Furthermore, we also showed that the homeostatic machinery in itself can affect the firing patterns of a neuron, as the afterhyperpolarizations seen in Fig 6A-D were concentration-dependent effects evoked by the electrogenic 3Na^+^/2K^+^ pump.

As the edNEG model was constructed by expanding a previous model [17] by (i) adding a glial domain and (ii) accounting for cellular swelling due to osmotic gradients, we put an extra emphasis on exploring how the presence of glia affected neuronal firing and swelling. In Fig 6, we showed that the glial support increased the tolerance level for neuronal firing and that the neuron could maintain steady-state firing for at least 90 s at frequencies up to 7.6 Hz in the presence of glia, but only up to 4.7 Hz when the glia domain was inactivated. Furthermore, the presence of glia reduced the swelling of the neuron from a maximum value of 3.7 % to a maximum value of 1.6 % during steady-state firing, and from 22.3 % to 16.3 % during depolarization block. The maximal neuronal and glial swelling coincided with a corresponding shrinkage of the extracellular space by 58.8 % of the original value for the version with glia, and 44.5 % for the version without glia. These quantitative predictions do of course depend on the included neuronal and glial mechanisms, the volume fractions, and the (sealed) boundary conditions used in the current simulation. However, they are in agreement with experimental studies, where reports of ECS shrinkage during SD range from 40 % to 78 % [20, 50–54].

Although the current implementation of the edNEG model contained only two neuronal compartments, the framework it was based upon can essentially be seen as a general framework for combining multicompartmental neural modeling with electrodiffusive ion concentration dynamics in neuroglial brain tissue. To our knowledge, the edNEG model is the first model to do this in a biophysically consistent manner, although many previous models have parts of the same functionality [13–16, 44, 47, 55–83].

### The outlook for an improved model of spreading depression

A key motivation for developing the edNEG model was its potential use in addressing SD and other pathological conditions associated with dramatic extracellular ion concentration changes.

SD was first described by Leão as a wave of silence propagating across cortex [84]. The spread of the wave coincides with shifts in the ECS K^+^ concentration by several tens of millimolar, DC-like voltage shifts in the ECS that may be as large as several tens of millivolts, swelling of neurons and glial cells, and changes in numerous other variables including the extracellular glutamate concentration and intracellular calcium concentrations [18, 23, 85]. The leading hypothesis, proposed by Grafstein already in 1956, is that diffusion of K^+^ through the ECS is the main propagator of the wave [25, 26]. However, buffering of K^+^ through the glial syncitium [86, 87], and electrical drift of K^+^ along the large DC-like voltage shifts [28, 88] are also likely to contribute to the wave propagation. Initiation of SD, and the leading edge of the SD wave, are often seen to occur in superficial (dendritic) layers of cortex and hippocampus, suggesting that dendritic membrane mechanisms play an important role for its pathophysiology [19, 20, 24, 30–32].

To our knowledge, only one computational model exists that has combined spatial propagation of SD with morphologically detailed neuron models [82]. However, this model was not based on an electrodiffusive formalism, and did not account for effects of extracellular potentials on neurodynamics and K^+^ transport. Other spatial models of SD [27–29, 40, 88] have been inspired by the coarse-grained *bi-domain* model [89], which previously has been used to simulate cardiac tissue [90, 91]. These models are electrodiffusive, and treat brain tissue as a homogeneous, coarse-grained, continuum, making them computationally efficient to allow for large scale simulations of SD propagation. However, they are limited in terms of neuronal detail, as none of them include fast neuronal mechanisms for action potential generation, or account for any morphological aspects of neurons, i.e., they do not account for the differences between dendritic and somatic layers.

The edNEG model was highly inspired by the previous tri-domain continuum model by Tuttle et al. 2019 [27], which is the most advanced of the spatial SD models. It includes neurons, glia, and extracellular space (Fig 7A), and it accounts for cellular swelling, a number of (slow) membrane mechanisms, and electrodiffusive ion concentration dynamics. Unlike other local models of ion concentration dynamics in tissue (see, e.g., [13, 44, 60, 71]), the edNEG model was based on the same kind of electrodiffusive formalism as the model by Tuttle et al. 2019 [27], and should in that regard be compatible with the tri-domain continuum framework used there (Fig 7A). We envision that the edNEG model can be integrated with this framework to obtain a *tri-times-two-domain* model (Fig 7B) that expands the functionality of the original tri-domain framework by accounting for (i) fast neuronal dynamics of AP firing and dendritic Ca^2+^ spiking, and (ii) differences between a deep (somatic) and superficial (dendritic) layer.

**Fig 7.**
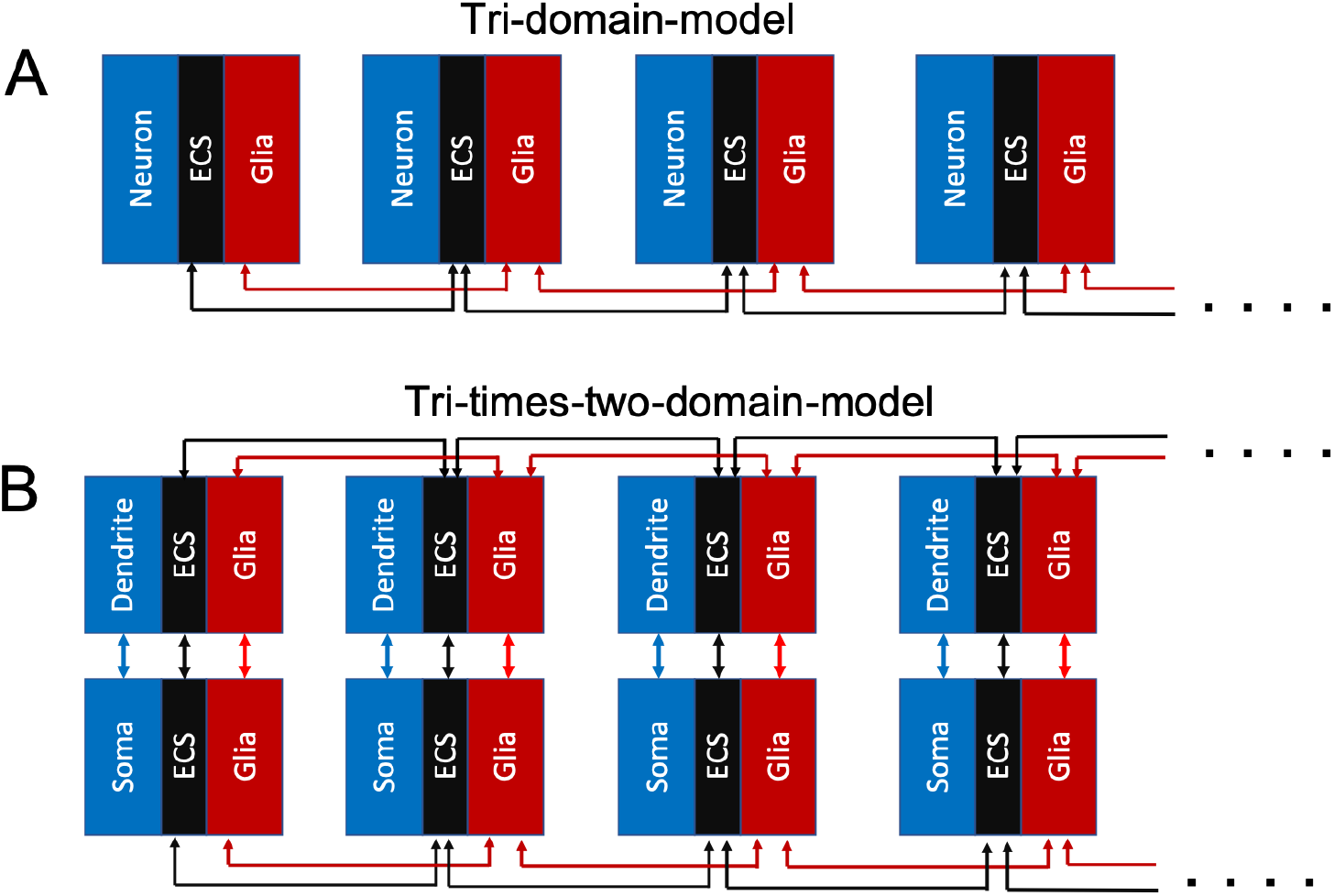
Tri-domain models for tissue dynamics. **(A)** Tri-domain model with a neuronal, ECS, and glial domain. At each point in space, a set of variables (voltage, ion concentrations, volume fractions) are defined for each of the three domains. At each local point (tricolored module), the three domains interact via transmembrane fluxes. In addition, modules interact with neighboring modules via spatiotemporal electrodiffusion in the glial domain (spatial buffering through glial syncytium), and the ECS domain. Neurons are typically assumed not to interact laterally in such a spatially continuous fashion. The spatial dynamics can, in principle, occur in all directions (3D), but a 1D illustration was used in the figure. **(B)** Tri-times-two domain model, where the local module has been replaced with the edNEG model so that it has two layers: a somatic (deep) layer, and a dendritic (superficial) layer. The local module then contains six domains: a neuronal, ECS, and glial domain in each of the two layers. Within each layer (*xy*-plane), the dynamics are modeled in the same fashion as in **(A)**. In addition, the tri-times-two domain model contains between-layer dynamics in terms of electrodiffusive transports intracellularly in neurons, intracellularly in the glial syncytium, and extracellularly. The key novelty is that the tri-times-two domain model can include different ion channels in the soma versus dendrites of neurons, and can capture somatodendritic signaling in neurons.

If the edNEG model is embedded in a tri-times-two domain framework for SD-susceptible brain tissue, it will become the first SD model that combines morphologically and biophysically detailed description of neurons with electrodiffusive continuum modeling of ion concentration- and voltage dynamics in neuron-ECS-glial-brain tissue at a large spatial scale. We anticipate that such a framework will be of great value for the field of neuroscience, partly because it gives a deepened insight into the balance between neuronal firing, ion homeostasis, and glial buffering at a local scale, partly because it may lead to new insight in the physiology of brain tissue in general, but most importantly because it invites detailed mechanistic studies of a number of pathological conditions associated with shifts in extracellular concentrations, such as SD, ischemic or hemorrhagic stroke, traumatic brain injury, migraine, and epileptic seizures [18, 20–23, 85, 92].

As a preliminary result, using the increase in the ECS K^+^-concentration as an indicator of SD initiation, the edNEG model was in agreement with the notion of earlier SD initiation in the dendritic layer, although layer differences were admittedly quite small in our simulations (Fig 5). We note, however, that the model was not in any way tuned to reproduce explicit SD data, and that these layer specificities followed directly from adopting a set of membrane mechanisms from a previous CA3 neuron model [33]. Putatively, larger differences between the layers could be obtained by adjusting either the somatic and dendritic ion channel conductances or the compartment sizes. In the current version, the neuronal soma and dendrite compartment were implemented with identical volumes and surface areas (cf. the version of the Pinsky-Rinzel model presented in [45]). If, for example, the neuronal surface-to-volume area was assumed to be larger in the dendritic layer, which would sound like a plausible assumption, we would expect a faster dissipation of the concentration gradients, and thus a more rapid increase in the ECS K^+^ concentration in the dendritic layer. Hence, if embedded in a tri-times-two continuum model for SD, there would be several approaches to retuning the edNEG model to make it in agreement with specific experimental data.

## Methods

### The Kirchoff-Nernst-Planck (KNP) framework for a tri-times-two compartment model

In a previous study [17], we derived the Kirchhoff-Nernst-Planck (KNP) framework [16, 38, 83, 93] for a closed-boundary system containing 2 × 2 compartments, representing a soma, a dendrite, and extracellular space (ECS) outside the soma and dendrite. In the edNEG model, we expanded the KNP framework to also include a glial domain, and to account for osmotically induced volume changes. The three domains (neuron + ECS + glia) were all represented as two-compartment models (one compartment in the soma layer and one in the dendrite layer). Within each layer, the neuron and glial domain interacted with the ECS through transmembrane currents (Fig 1). Volume changes were due to osmotic pressure gradients that we implemented as functions of the ionic concentrations (see section titled Volume dynamics). Geometrical parameters, including initial volumes, are listed in Table 1.

**Table 1.** Geometrical parameters

| Parameter | Value | Reference |
| --- | --- | --- |
| $\Delta x$ (distance between the two layers) | $667 \cdot 10^{-6} \text{ m}$ | [17] |
| $A_m$ (membrane area of each cellular compartment) | $616 \cdot 10^{-12} \text{ m}^2$ | [17] |
| $A_i$ (intracellular cross-section areas) | $\alpha \cdot A_m^\dagger$ | [17] |
| $A_e$ (extracellular cross-section area) | $A_i/2$ | [17] |
| $V_{sn,0}, V_{dn,0}$ (initial neuronal volumes) | $1437 \cdot 10^{-18} \text{ m}^3$ | [17] |
| $V_{se,0}, V_{de,0}$ (initial extracellular volumes) | $718.5 \cdot 10^{-18} \text{ m}^3$ | [17] |
| $V_{sg,0}, V_{dg,0}$ (initial glial volumes) | $1437 \cdot 10^{-18} \text{ m}^3$ | |

<sup>†</sup> The parameter $\alpha$ determines the coupling strength between the soma and dendrite layers of the model. Its default value was 2, same as in [17].

#### Electrodiffusion

Two kinds of fluxes transport ions in the system: transmembrane fluxes and axial fluxes. The axial fluxes are driven by electrodiffusion, and we describe them using the Nernst-Planck equation. It follows that the intracellular flux density of the neuron for ion species *k* is expressed as:

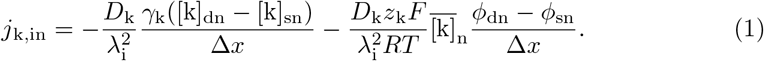

In Eq 1, we use the same notation as in [17] so that *D*_k_ is the diffusion constant, *γ*_k_ is the fraction of freely moving ions, that is, ions that are not buffered or taken up by the endoplasmatic reticulum, *λ*_i_ is the tortuosity, which represents hindrances in free diffusion due to obstacles, *γ*_k_([k]_dn_ − [k]_sn_)/Δ*x* is the longitudinal concentration gradient, *z*_k_ is the charge number of ion species *k*, *F* is the Faraday constant, *R* is the gas constant, *T* is the absolute temperature, 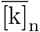 is the average concentration, that is, *γ*_k_([k]_dn_ + [k]_sn_)/2, and (*ϕ*_dn_ − *ϕ*_sn_)/Δ*x* is the longitudinal potential gradient. Likewise, the extracellular flux densities and the glial intracellular flux densities are described, respectively, by

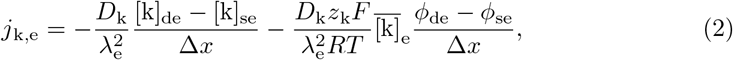

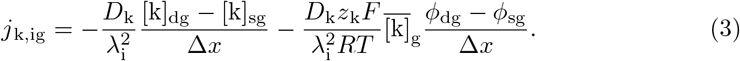

All ions can move freely in the extracellular and glial space, thus, *γ*_k_ is not included in Eqs 2 and 3. Diffusion constants, tortuosities, and intracellular fractions of freely moving ions are listed in Table 2.

**Table 2.** Diffusion constants, tortuosities, and intraneuronal fractions of freely moving ions^†^

| Parameter | Value | Reference |
| --- | --- | --- |
| $D_{\text{Na}}$ ($\text{Na}^+$ diffusion constant) | $1.33 \cdot 10^{-9} \text{ m}^2/\text{s}$ | [93, 94] |
| $D_{\text{K}}$ ($\text{K}^+$ diffusion constant) | $1.96 \cdot 10^{-9} \text{ m}^2/\text{s}$ | [93, 94] |
| $D_{\text{Cl}}$ ($\text{Cl}^-$ diffusion constant) | $2.03 \cdot 10^{-9} \text{ m}^2/\text{s}$ | [93, 94] |
| $D_{\text{Ca}}$ ($\text{Ca}^{2+}$ diffusion constant) | $0.71 \cdot 10^{-9} \text{ m}^2/\text{s}$ | [93, 94] |
| $\lambda_i$ (intracellular tortuosity) | 3.2 | [36, 38] |
| $\lambda_e$ (extracellular tortuosity) | 1.6 | [36, 38] |
| $\gamma_{\text{Na}}, \gamma_{\text{K}}, \gamma_{\text{Cl}}$ (intraneuronal fractions of free ions) | 1 | [17] |
| $\gamma_{\text{Ca}}$ (intraneuronal fraction of free ions) | 0.01 | [17] |

<sup>†</sup> The table is adopted from [17].

**Table 3.** Temperature and physical constants^†^

| Parameter | Value | Reference |
| --- | --- | --- |
| $T$ (absolute temperature) | 309.14 K | [17, 44] |
| $F$ (Faraday constant) | $9.648 \cdot 10^4$ C/mol | |
| $R$ (gas constant) | 8.314 J/(mol K) | |

<sup>†</sup> This table is adopted from [17].

#### Ion conservation

To keep track of all ions in the system, we solve six differential equations for each ion species *k*. Conservation of ions gives:

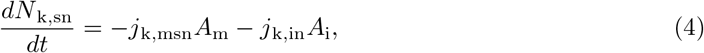

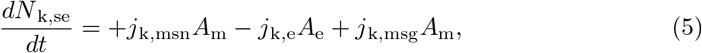

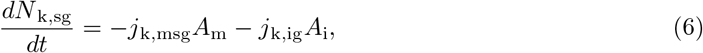

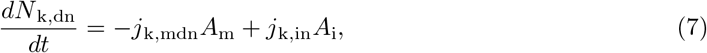

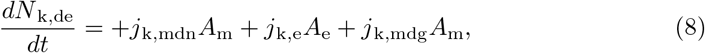

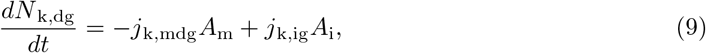

where *N* is the amount of substance, in units of mol. To find the change in *N*, all ion flux densities are multiplied by the area they go through. The variable *j*_k,m_ represents the sum of all membrane flux densities of ion species *k*, and *j*_k,in_, *j*_k,e_, and *j*_k,ig_ represent the axial flux densities. To find the ion concentrations, which appear in many of the following equations, we divide the amounts of substance in a compartment by the compartment volume at the beginning of each time step.

We insert the Nernst-Planck equation for the axial flux density (Eq 1) into Eq 4 and get:

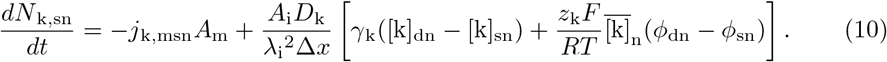

In Eq 10, [k]_dn_ and [k]_sn_ are the intraneuronal ion concentrations of the dendrite and soma, defined as *N* _k,dn_*/V* _dn_ and *N* _k,sn_*/V* _sn_, respectively. We define the voltage variables *ϕ*_dn_ and *ϕ*_sn_ below.

#### Six constraints to derive *ϕ*

If we have four ion species (Na^+^, K^+^, Cl^−^, and Ca^2+^) in six compartments, we get 24 equations to solve (Eqs 4-9 times four) and 30 unknowns (*N* and *ϕ*). We overcome this by defining *ϕ* in terms of ion concentrations using a set of constraints similar to those used in [17].

1. *Arbitrary reference point for ϕ.* The first constraint is simple; we can choose an arbitrary reference point for *ϕ*. We define it to be outside the dendrite, which gives us:

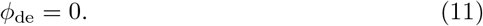
2. *Neuronal membrane is a parallel plate capacitor (dendrite).* As the second constraint, we use that the membrane is a parallel plate capacitor. This means that it will always separate a charge *Q* on one side from an opposite charge −*Q* on the other side. This gives rise to a voltage difference across the membrane

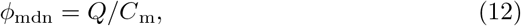

where *C*_m_ is the total capacitance of the membrane, i.e., *C*_m_ = *c*_m_*A*_m_, where *c*_m_ is the capacitance per membrane area. We know, by definition, that *ϕ*_mdn_ = *ϕ*_dn_ *− ϕ*_de_, and since *ϕ*_de_ = 0, we get:

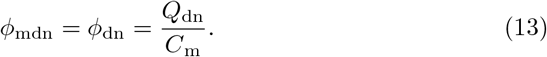 We assume bulk electroneutrality, meaning that all net charge in the dendritic compartment must be on the membrane. It follows that 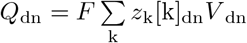, where *F* is the Faraday constant, *z*_k_ is the charge number of ion species *k*, [k]_dn_ is the ion concentration, and *V* _dn_ is the volume. By inserting this into Eq 13, we get

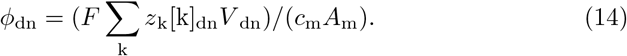
3. *Neuronal membrane is a parallel plate capacitor (soma).* The second constraint also applies to the soma, and gives us the criterion:

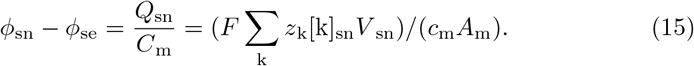 Here, the outside potential is not set to zero, so this constraint is not sufficient to determine *ϕ*_sn_ and *ϕ*_se_ separately.
4. *Glial membrane is a parallel plate capacitor (dendrite layer).* The glial membrane is no different than the neuronal membrane when it comes to acting as a parallel plate capacitor, so we get:

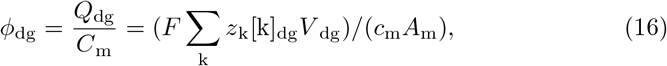

where we have used that *ϕ*_de_ = 0.
5. *Glial membrane is a parallel plate capacitor (soma layer).* Constraint number (iv) also applies to the soma layer, and gives us:

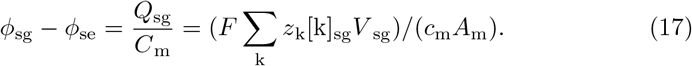 We can now calculate *ϕ*_dn_ and *ϕ*_dg_ from Eqs 14 and 16 but to determine *ϕ*_sn_, *ϕ*_se_, and *ϕ*_sg_, we need a sixth constraint.
6. *Current anti-symmetry.* The sixth constraint is to ensure charge (anti-)symmetry. We must define the initial conditions so that the membrane separates a charge *Q* on one side from an opposite charge −*Q* on the other side, and the system dynamics so that it stays this way. The membrane fluxes (alone) fulfill this criterion, since a charge that leaves a compartment automatically pops up on the other side of the membrane, making sure that *dQ*_i_*/dt* = −*dQ*_e_*/dt*. For the axial fluxes to fullfill the criterion, we must have that:

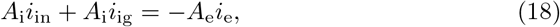

where *i* stands for current density. We find expressions for *i*_in_, *i*_ig_, and *i*_e_, by multiplying Eqs 1-3 by *Fz*_k_ and sum over all ion species *k*. Expressions for the current densities then become:

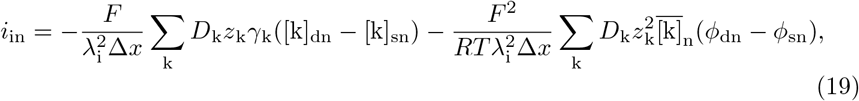

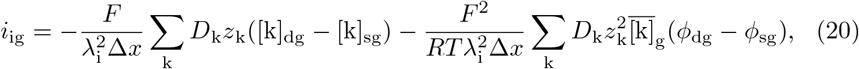

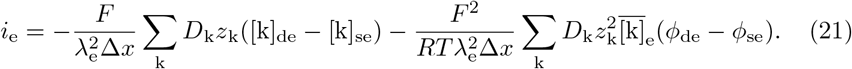

The first term in Eq 19 is the diffusion current density and is defined by the ion concentrations:

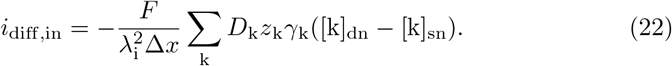

The second term is the field driven current density

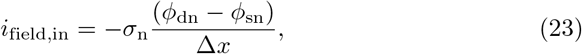

where *σ*_n_ is the conductivity:

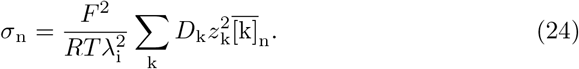

Likewise, Eq 20 can be written in terms of *i*_diff,ig_, *i*_field,ig_, and *σ*_g_, and Eq 21 in terms of *i*_diff,e_, *i*_field,e_, and *σ*_e_. We combine Eqs 18-21 and obtain:

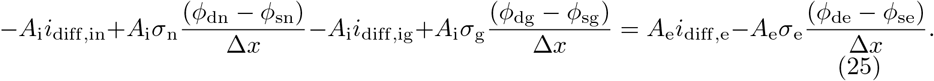

In Eq 25, we know *ϕ*_dn_, *ϕ*_de_, and *ϕ*_dg_ from Eqs 14, 11, and 16, and *i*_diff_ and *σ* from the ion concentrations. We solve Eqs 15, 17, and 25 to find *ϕ*_sn_, *ϕ*_sg_, and *ϕ*_se_:

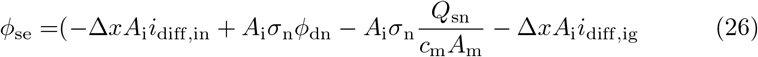

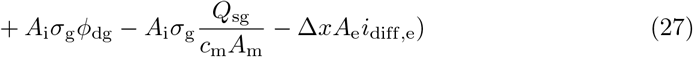

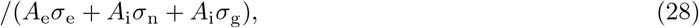

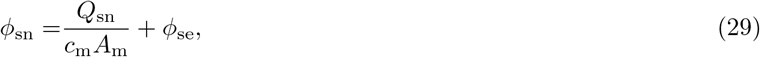

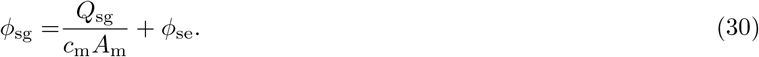

### Neuronal membrane mechanisms

The neuronal membrane mechanisms were equal to those in [17]. We list them again here for easy reference.

#### Leakage channels

Both neuronal compartments contained Na^+^, K^+^, and Cl^−^ leak currents. The flux densities were modeled as in [44]:

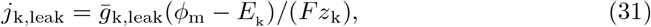

where *k* denotes the ion species, 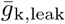 is the ion conductance, *ϕ*_m_ is the membrane potential, *E*_k_ is the reversal potential, *F* is the Faraday constant, and *z*_k_ is the charge number. Reversal potentials are given by the Nernst equation:

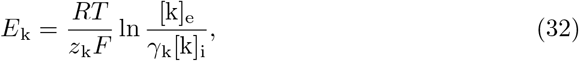

where *R* is the gas constant, *T* is the absolute temperature, *γ*_k_ is the intracellular fraction of freely moving ions, and [k]_e_ and [k]_i_ are the extra- and intracellular concentrations of ion species *k*, respectively.

#### Active ion channels

We adopted all active ion channels from the Pinsky-Rinzel model [33]. This included Na^+^ and K^+^ delayed rectifier fluxes in the soma (*j*_Na_, *j*_DR_), and a voltage-dependent Ca^2+^ flux (*j*_Ca_), a voltage-dependent K^+^ afterhyperpolarization flux (*j*_AHP_), and a Ca^2+^-dependent K^+^ flux (*j*_C_) in the dendrite:

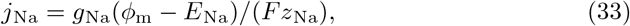

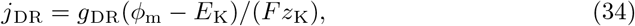

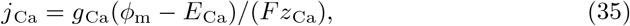

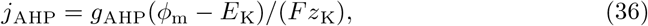

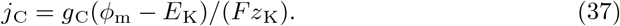

Here, *g*_k_ is the ion conductance, *ϕ*_m_ is the membrane potential, *E*_k_ is the reversal potential, *F* is the Faraday constant, and *z*_k_ is the charge number for ion species *k*. We used the Hodkin-Huxley formalism to model the voltage-dependent conductances, with differential equations for the gating variables:

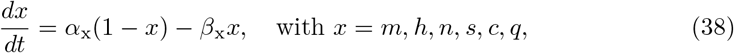

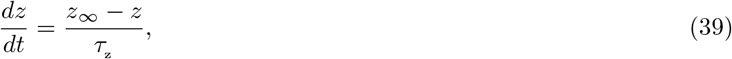

and

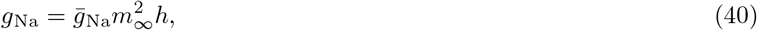

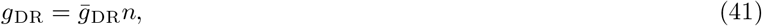

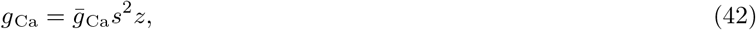

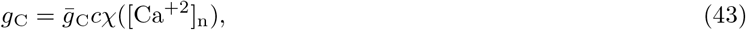

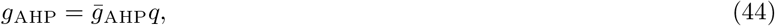

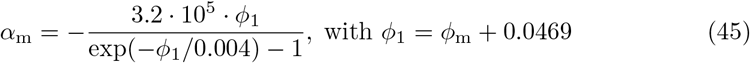

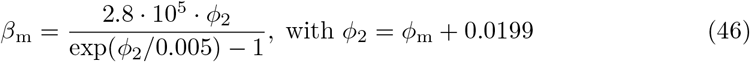

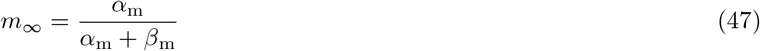

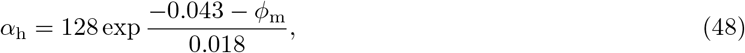

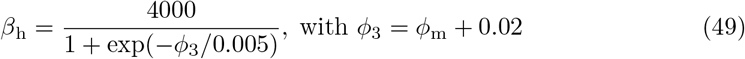

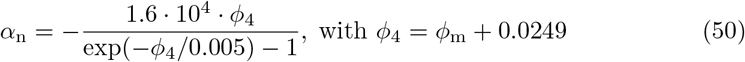

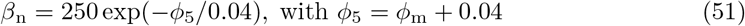

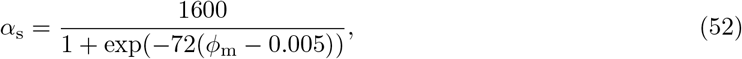

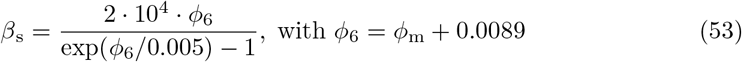

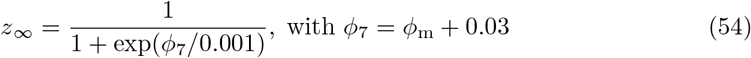

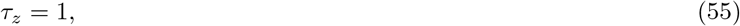

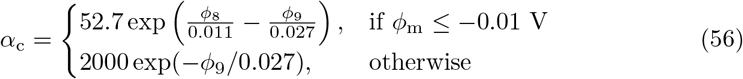

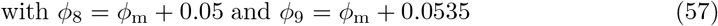

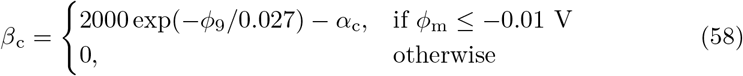

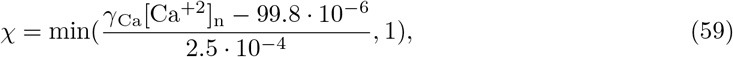

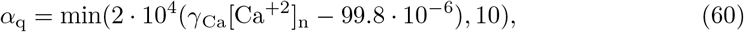

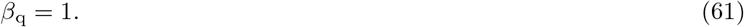

In Eqs 38-61, rates (*α*’s, *β*’s) are in units of 1/s, *τ* _z_ is in units of s, and voltages *ϕ* are in units of V.

#### Homeostatic mechanisms

Both neuronal compartments contained a 3Na^+^/2K^+^ pump, a K^+^/Cl^−^ cotransporter (KCC2), a Na^+^/K^+^/2Cl^−^ cotransporter (NKCC1), and a 2Na^+^/Ca^2+^ exchanger. The functional forms of the pumps and cotransporters were taken from [44], while the 2Na^+^/Ca^2+^ exchanger was modeled as in [17]:

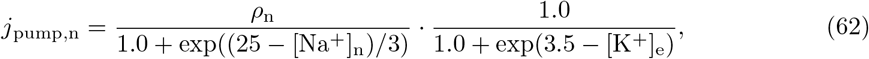

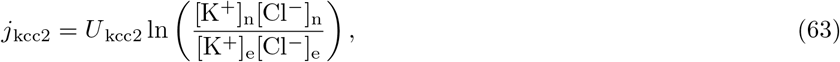

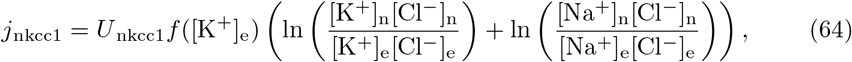

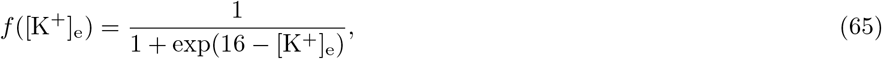

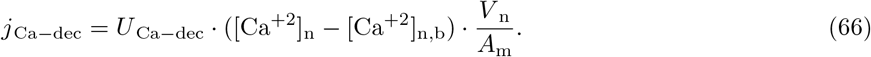

Here, *ρ*_n_, *U* _kcc2_, and *U* _nkcc1_ are pump and cotransporter strengths, *U* _Ca−dec_ is the Ca^2+^ decay rate, and [Ca^+2^]_n,b_ is the basal Ca^2+^ concentration..

### Glial membrane mechanisms

We modeled the glia as an astrocytic domain and adopted the membrane mechanisms from [38]. They included Na^+^ and Cl^−^ leak channels, modeled as in Eq 31, an inward rectifying K^+^ channel, and a 3Na^+^/2K^+^ pump:

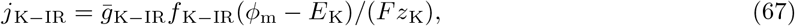

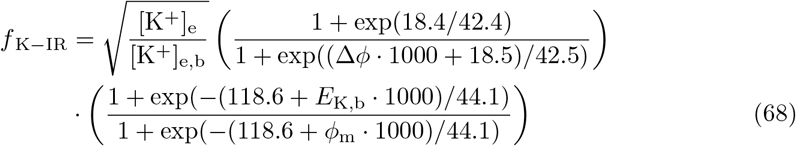

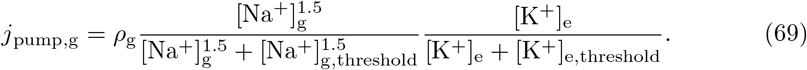

Here, 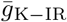 is the K^+^ ion conductance, *ϕ*_m_ is the membrane potential, *E*_K_ is the K^+^ reversal potential, *F* is the Faraday constant, *z*_K_ is the K^+^ charge number, [K^+^]_e,b_ is the basal K^+^ concentration in the extracellular space, 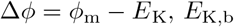 is the reversal potential for K^+^ at basal concentrations, *ρ*_g_ is the pump strength, and [Na^+^]_g,threshold_ and [K^+^]_e,threshold_ are the pump’s threshold concentrations for Na^+^ and K^+^, respectively. We included the same set of membrane mechanisms in both astrocytic compartments.

### Volume dynamics

To calculate the osmotically induced volume changes *dV/dt*, we used the formalism outlined in [42]:

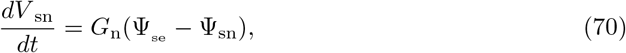

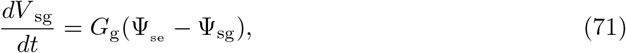

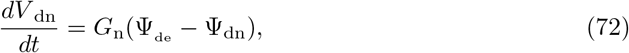

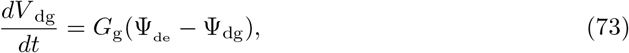

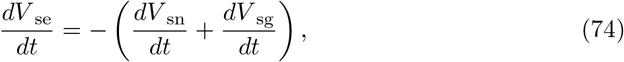

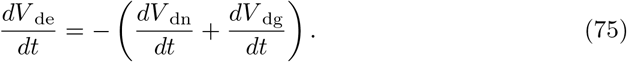

Here, *G*_n_ and *G*_g_ are the neuronal and glial water permeabilities, respectively, given in units of m^3^/Pa/s, and ψ is the water potential, given in units of Pa. We assumed the hydrostatic pressure differences to be zero, so that water flow was driven by osmotic pressure differences only, and we calculated the solute potentials from:

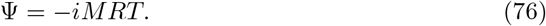

Here, *i* is the ionization factor (van’t Hoff factor), which is 1 for ions, *M* is the osmotic concentration of solutes measured in moles per cubic meter, *R* is the gas constant, and *T* is the absolute temperature. Equations 74 and 75 follow from the assumption that the total volume did not change, that is, the system was closed.

Like in [27], we only considered effects of transmembrane water flow, and intra-domain water flow due to hydrostatic pressures were neglected. As predicted in a previous study, bulk flow at physiological hydrostatic pressure is expected to be low [95].

### Model summary

To keep track of all ions in the system, we solved six differential equations for each ion species *k*:

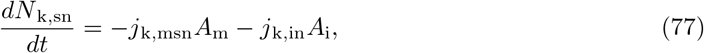

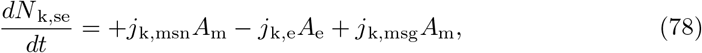

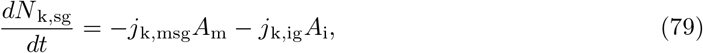

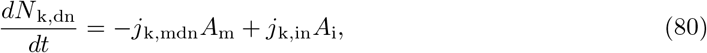

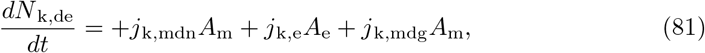

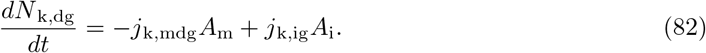

The total membrane flux densities are summarized here:

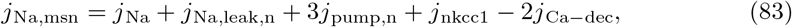

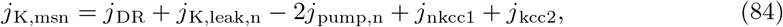

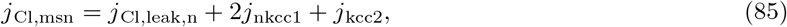

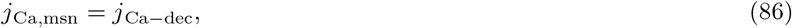

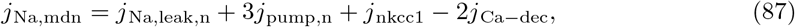

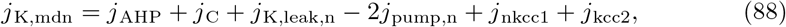

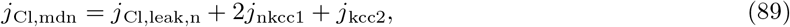

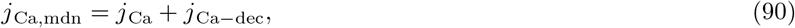

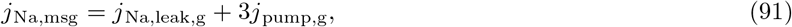

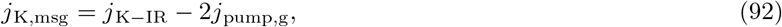

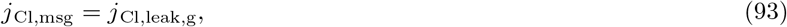

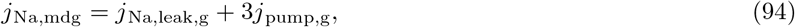

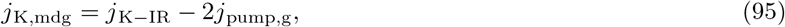

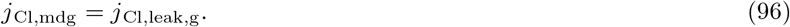

At each time step, we derived *ϕ* algebraically in all six compartments:

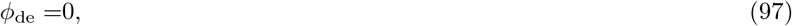

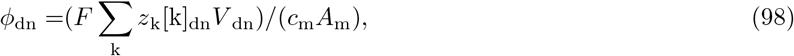

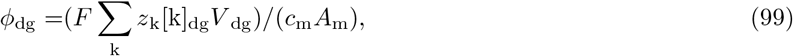

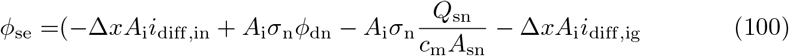

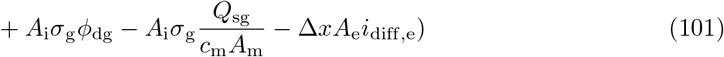

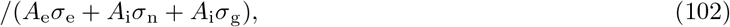

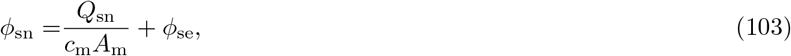

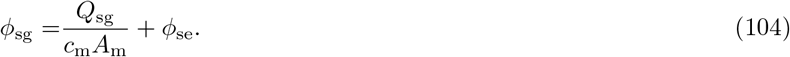

Volume dynamics was given by:

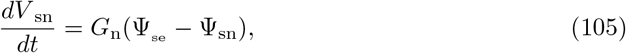

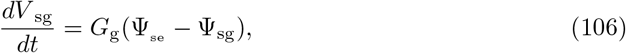

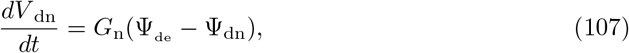

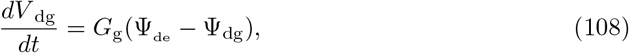

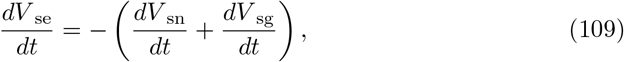

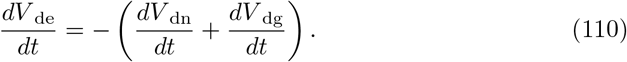

Fig 1 summarizes the model and model parameters are listed in Tables 1-5.

## Simulations

### Model tuning

The edNEG model combined two previous models, one consisting of a neuron and ECS [17], and the other of a glial domain (astrocyte) and ECS [38]. When we combined the models, we set the initial ionic concentrations in the neuron identical to those in [17], the initial ionic concentrations in the glial domain identical to those in [38], and made the two cells share the same ECS where we set the initial concentrations equal to those in the previous glia model [38]. As these initial concentrations (Table 5) differed from the initial ECS concentrations in the previous neuron model [17], the neuron was not in equilibrium with the (new) ECS. This was because the altered ECS concentrations gave rise to altered concentration-dependent activity of the ion pumps, cotransporters, and ionic currents through ion channels. We found that the leakage currents were most important, and that a re-tuning of the leak conductances (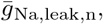 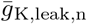, and 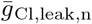) in the neuron model was sufficient to obtain a system with a plausible resting state. The tuning was done by requiring that the initial leakage currents should be identical to those in [17], i.e., we set:

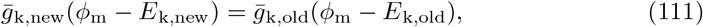

with *ϕ*_m_ being the resting potential in [17] (−67.7 mV), *E*_k,old_ being the reversal potential for ion species *k* at steady state in [17], and *E*_k,new_ being the reversal potential obtained by the new initial ion concentrations (Table 5). By solving this equation, we obtained a final set of passive conductances for the neuronal membrane (Table 4). After calibrating (running it for 5000 s) the edNEG model with the (new) derived passive conductances, it settled at a resting state where the neuronal resting membrane potential was −70.3 mV, and the glial membrane resting potential was −82.6 mV, which was close to the original resting potentials for the neuron and glial domain (original values were −67.7 mV [17] and −83.6 mV [38], respectively). The reason why the new and original values were not identical was that not only the leakage currents, but also the ion pumps and cotransporters, and to a small extent the active channels, were active at rest.

**Table 4.** Membrane parameters

| Parameter | Value | Reference |
| --- | --- | --- |
| $c_m$ | $3 \cdot 10^{-2}$ F/m<sup>2</sup> | [33, 45] |
| $\bar{g}_{Na,leak,n}$ | 0.246 S/m<sup>2</sup> | eq. 111 |
| $\bar{g}_{K,leak,n}$ | 0.245 S/m<sup>2</sup> | eq. 111 |
| $\bar{g}_{Cl,leak,n}$ | 0.668 S/m<sup>2</sup> | eq. 111 |
| $\bar{g}_{Na}$ | 300 S/m<sup>2</sup> | [33, 45] |
| $\bar{g}_{DR}$ | 150 S/m<sup>2</sup> | [33, 45] |
| $\bar{g}_{Ca}$ | 141 S/m<sup>2</sup> | tuned |
| $\bar{g}_{AHP}$ | 8 S/m<sup>2</sup> | [33, 45] |
| $\bar{g}_C$ | 150 S/m<sup>2</sup> | [33, 45] |
| $\rho_n$ | $1.87 \cdot 10^{-6}$ mol/(m<sup>2</sup>s) | [17, 44] |
| $U_{kcc2}$ | $7.0 \cdot 10^{-7}$ mol/(m<sup>2</sup>s) | [17, 44] |
| $U_{nkcc1}$ | $2.33 \cdot 10^{-7}$ mol/(m<sup>2</sup>s) | [17, 44] |
| $U_{Ca-dec}$ | 75 s<sup>-1</sup> | [33, 45] |
| $\bar{g}_{Na,leak,g}$ | 1 S/m<sup>2</sup> | [38] |
| $\bar{g}_{Cl,leak,g}$ | 0.5 S/m<sup>2</sup> | [38] |
| $\bar{g}_{K-IR}$ | 16.96 S/m<sup>2</sup> | [38] |
| $\rho_g$ | $1.12 \cdot 10^{-6}$ mol/(m<sup>2</sup>s) | [38] |
| $[Na^+]_{g,threshold}$ | 10 mM | [38] |
| $[K^+]_{e,threshold}$ | 1.5 mM | [38] |
| $G_n$ | $2 \cdot 10^{-23}$ m<sup>3</sup>/Pa/s | [96] |
| $G_g$ | $5 \cdot 10^{-23}$ m<sup>3</sup>/Pa/s | [41] |

**Table 5.** Initial conditions

| Variables | Pre-calibrated | Post-calibrated<sup>1</sup> | Reference |
| --- | --- | --- | --- |
| $\phi_{mn,0}^\dagger$ | -67.7 mV | -70.3 mV | [17] |
| $\phi_{mg,0}^\dagger$ | -83.6 mV | -82.6 mV | [38] |
| $[Na^+]_{n,0}$ | 16.9 mM | 18.4 mM | [17] |
| $[Na^+]_{e,0}$ | 144.622 mM | 144.0 mM | [38] |
| $[Na^+]_{g,0}$ | 15.189 mM | 14.0 mM | [38] |
| $[K^+]_{n,0}$ | 139.5 mM | 137.1 mM | [17] |
| $[K^+]_{e,0}$ | 3.082 mM | 3.8 mM | [38] |
| $[K^+]_{g,0}$ | 99.959 mM | 102.0 mM | [38] |
| $[Cl^-]_{n,0}$ | 5.4 mM | 4.5 mM | [17] |
| $[Cl^-]_{e,0}$ | 133.71 mM | 133.7 mM | [38] |
| $[Cl^-]_{g,0}$ | 5.145 mM | 6.0 mM | [38] |
| $[Ca^{2+}]_{n,0}$ | 0.01 mM* | 0.01 mM* | [17] |
| $[Ca^{2+}]_{e,0}$ | 1.1 mM | 1.1 mM | [17] |
| $n_0$ | 0.0003 | 0.0002 | [17] |
| $h_0$ | 0.999 | 0.9997 | [17] |
| $s_0$ | 0.007 | 0.0057 | [17] |
| $c_0$ | 0.005 | 0.0042 | [17] |
| $q_0$ | 0.011 | 0.0092 | [17] |
| $z_0$ | 1.0 | 1.0 | [17] |

<sup>1</sup> Values with more decimals included were read to/from file and used in the simulations. (Available at https://github.com/CINPLA/edNEGmodel_analysis.)
<sup>†</sup> $\phi_m$ is not an independent state variable, but defined at each time point from the ion concentrations.
\* Only 1% of the total intracellular $Ca^{2+}$ , that is, a 100 nM, was assumed to be free (unbuffered).

To obtain comparable spike shapes between the edNEG model and the original Pinsky-Rinzel model (Fig 2), we manually tuned the Ca^2+^ conductance of the neuron 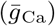, as well as the coupling strength between the soma and dendrite layers. The coupling strength was regulated by adjusting the intracellular cross-section areas *αA*_i_ (cf. Table 1) by adjusting the unit-less parameter *α*. The parameter *α* was 2 for simulations with *stong* coupling between the soma and dendrite layers, and 0.51 for simulations with weak coupling. Strong coupling was used in all simulations except in the simulation shown in Fig 2C where it was 0.51. The Ca^2+^ conductance 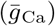, along with the other membrane mechanisms, were the same in all simulations (values as in Table 4).

### Initial conditions

Before tuning the edNEG model, we defined its initial volumes (Table 1), amounts of ions, membrane potentials, and gating variables (Table 5, Pre-calibrated column) using values from the two previous models in [17] and [38]. After re-tuning selected parameters (as described in the previous subsection), the system was close to, but not strictly in equilibrium, and for this reason we calibrated the edNEG model for 5000 s. The water permeabilities were set to zero during the calibration.

We wrote the final values from the calibration to file (see Table 5, Post-calibrated column) and used them as initial conditions in all simulations shown throughout this paper. Note that the edNEG model takes amounts of ions (in units of mol) as input, while we have listed ion concentrations in Table 5. The post-calibrated values of the ion concentrations correspond to the following reversal potentials: *E*_Na,n_ = 55 mV, *E*_Na,g_ = 62 mV, *E*_K,n_ = −96 mV, *E*_K,g_ = −88 mV, *E*_Cl,n_ = −90 mV, *E*_Cl,g_ = −83 mV, and *E*_Ca,n_ = 124 mV.

To ensure charge symmetry and electoneutrality, we defined a set of static residual charges, based on the initial amounts of ions. These represent negatively charged macromolecules present in real cells. We defined them as constant amounts of ion species X^−^ with charge number *z*_X_ = −1 and diffusion constant *D*_X_ = 0. To ensure strict electroneutrality, we did not read residual charges to/from file, but calculated them at the beginning of each simulation.They were given by the following expressions:

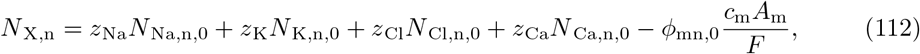

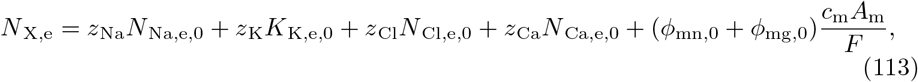

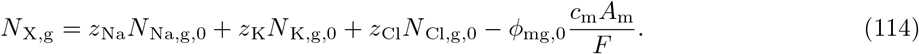

Additionally, we introduced a set of static residual molecules to ensure zero osmotic pressure gradients across the membranes at the beginning of each simulation. These were defined as osmotic concentrations of a molecule M:

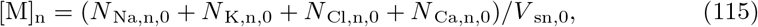

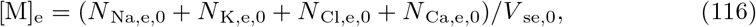

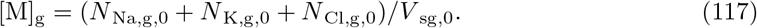

### Stimulus current

We stimulated the neuron like we did in [17], that is, by applying a K^+^ injection current *i*_stim_ into the soma, and removing the same amount of K^+^ from the corresponding extracellular compartment to ensure ion conservation:

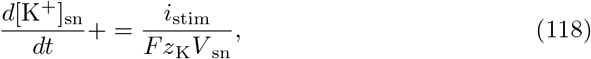

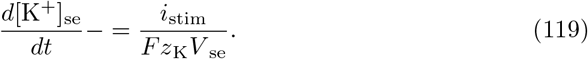

### Numerical implementation

We implemented the code in Python 3.6 and solved the differential equations using the solve ivp function from SciPy with its Runge-Kutta method of order 3(2). We set the maximal allowed step size to 10^−4^. The code can be downloaded from https://github.com/CINPLA/edNEGmodel and https://github.com/CINPLA/edNEGmodel_analysis.

